# GABA and glycine synaptic release on axotomized motoneuron cell bodies promotes motor axon regeneration

**DOI:** 10.1101/2024.11.18.623863

**Authors:** Ryan L. Wood, Paula M. Calvo, William M. McCallum, Arthur W. English, Francisco J. Alvarez

## Abstract

Motor axon regeneration after traumatic nerve injuries is a slow process that adversely influences patient outcomes because muscle reinnervation delays result in irreversible muscle atrophy and suboptimal axon regeneration. This advocates for investigating methods to accelerate motor axon growth. Electrical nerve stimulation and exercise both enhance motor axon regeneration in rodents and patients, but these interventions cannot always be easily implemented. A roadblock to uncover novel therapeutic approaches based on the effects of activity is the lack of understanding of the synaptic drives responsible for activity-mediated facilitation of axon regeneration. We hypothesized that the relevant excitatory inputs facilitating axon regrowth originate in GABA/glycine synapses which become depolarizing after downregulation of the potassium chloride cotransporter 2 in motoneurons following axotomy. To test this, we injected tetanus toxin (TeTx) in the tibialis anterior (TA) muscle of mice to block the release of GABA/glycine specifically on TA motoneurons. Thereafter, we axotomized all sciatic motoneurons by nerve crush and analyzed the time-courses of muscle reinnervation in TeTx- treated (TA) and untreated (lateral gastrocnemius, LG) motoneurons. Muscle reinnervation was slower in TA motoneurons with blocked GABA/glycine synapses, as measured by recovery of M-responses and anatomical reinnervation of neuromuscular junctions. Post-hoc immunohistochemistry confirmed the removal of the vesicular associated membrane proteins 1 and 2 by TeTx activity, specifically from inhibitory synapses. These proteins are necessary for exocytotic release of neurotransmitters. Therefore, we conclude that GABA/glycine neurotransmission on regenerating motoneurons facilitates axon growth and muscle reinnervation and discuss possible interventions to modulate these inputs on regenerating motoneurons.

## Introduction

The United States alone reports yearly over 200,000 nerve injuries resulting from physical trauma (Taylor *et al*., 2008), with the actual incidence likely being higher considering the diversity of presentations and comorbidities with other types of tissue damage. Nerve injury disconnects sensory and motor axons from their peripheral targets and although axons eventually regrow, functional recovery is frequently disappointing. For example, despite continuous advances in nerve-repair and microsurgical techniques, only 10% of adults recover near-normal function after median or ulnar nerve repair (Lundborg, 2003; Brushart, 2011). Thus, not only the incidence of nerve injuries is high, but patients often face life-long disabilities. To optimize therapies, it is critical to better understand regeneration mechanisms of motor axons. One often cited impediment for recovery is the slow speed of axon regeneration (1-3 mm per day). Regenerative capacity declines with time after injury (Fu & Gordon, 1995b; a; Furey *et al*., 2007) and prolonged muscle denervation induces irreversible muscle fiber atrophy and replacement with connective tissue and fat (Carlson, 2014). Thus, the speed at which muscles become reinnervated is one of the best predictors of recovery outcomes. Several interventions have been shown to accelerate regeneration (Gordon & Sulaiman, 2013; Scheib & Hoke, 2013; Gordon, 2016), but there is incomplete knowledge of endogenous mechanisms recruited by these interventions to promote axon growth. A deeper understanding of these mechanisms could be exploited in the future to improve outcomes in patients that undergo nerve surgeries..

Electrical nerve stimulation and exercise are two methods that recently have shown promise to enhance axon regeneration and motor function recovery after injury in experimental animals and also in limited clinical trials (English *et al*., 2007; Gordon *et al*., 2009; English *et al*., 2014; Gordon & English, 2016; Zuo *et al*., 2020). Both methods require motoneuron activation and BDNF-TrkB signaling (Al-Majed *et al*., 2000a; Al-Majed *et al*., 2000b; Al-Majed *et al*., 2004; English *et al*., 2007; Gordon *et al*., 2009; English *et al*., 2011; English *et al*., 2014; Gordon & English, 2016), but the synaptic drives that mediate activity-dependent regeneration in injured motoneurons remain unknown. We proposed that activity in GABAergic/glycinergic synapses on cell bodies of axotomized motoneurons might be a key contributor (Akhter, 2020). Such a mechanism offers new avenues to enhance motor axon regeneration, however it is contrary to prevailing interpretations of synaptic plasticity on axotomized motoneurons. After axotomy most excitatory synaptic boutons are removed from the cell body of injured motoneurons while there is partial retention of inhibitory synapses (Alvarez *et al*., 2011; Alvarez *et al*., 2020). This change in synaptic composition is usually explained as necessary for inducing a “quiet” phase, in which motoneurons are electrically silenced to focus resources on gene and protein expression specific to regeneration (Carlstedt & Cullheim, 2000; Oliveira *et al*., 2004; Navarro *et al*., 2007). However, this contradicts evidence showing that activity promotes regeneration. Moreover, “inhibitory” synapses lose inhibitory capacity on axotomized motoneurons because expression of the potassium-chloride co-transporter 2 gene (*kcc2)* is suppressed quickly after axotomy and this is followed by removal of membrane KCC2 protein soon after (Akhter *et al*., 2019). Loss of membrane KCC2 elevates intracellular chloride and result in GABA-mediated depolarizations and Ca^2+^ transients in brainstem motoneurons (Nabekura *et al*., 2002; Toyoda *et al*., 2003). These synaptic changes are reminiscent of earlier development in which low KCC2 expression, GABA depolarization and Ca^2+^ entry are all necessary for cell migration and neurite and axon elongation (Takayama & Inoue, 2004; Sernagor *et al*., 2010). This prompted us (Akhter, 2020), and others (Tatetsu *et al*., 2012), to suggest that GABA/glycine synaptic release on adult axotomized motoneurons in the absence of KCC2 might facilitate regenerative mechanisms.

To test this hypothesis, we used tetanus toxin (TeTx) to block inhibitory synapses specifically on one target pool of axotomized motoneurons and compared their regeneration to non-tetanized axotomized motoneurons. Motoneuron axotomy was obtained by a mid-thigh sciatic nerve crush (SNC), avoiding complexities due to errors in regeneration target specificity and allowing for faster regeneration and muscle reinnervation (Rotterman *et al*., 2024). TeTx injected in muscle retrogradely transports to motoneuron cell bodies where it jumps transynaptically into pre-motor neurons to block synaptic vesicle exocytosis and neurotransmitter release preferentially at GABA/glycine synapses through cleavage of vesicle-associated membrane proteins (VAMPs) (Brooks *et al*., 1957; Price *et al*., 1975; Schwab & Thoenen, 1978; Collingridge & Davies, 1982; Kanda & Takano, 1983; Schiavo *et al*., 1992; Schiavo *et al*., 2000; Rossetto *et al*., 2001; Lalli *et al*., 2003; Gonzalez-Forero *et al*., 2005; Fabris *et al*., 2022). To avoid systemic and off-target toxin effects we used very low doses of TeTx to transiently reduce GABA and/or glycine release specifically on one motor pool for a few weeks. The method was validated by quantifying *in vivo* reversible changes in motor unit excitability on tetanized motor pools and by observing VAMP protein reductions specifically in inhibitory synapses on these motor pools. After analyzing muscle reinnervation rates with functional and anatomical methods we found that TeTx-block of GABA and/or glycine release slowed muscle reinnervation suggesting these neurotransmitters exert axon growth promoting actions on regenerating motoneurons.

## Materials and Methods

### Ethical Standards

This study used wild type mice of both sexes. All animal care, procedures, and euthanasia were performed under prior approval by the Institutional Animal Care and Use Committee of Emory University (protocols PROTO202100174) and in accordance with the NIH Guide for the Care and Use of Laboratory Animals. Mice were housed based on sex in groups of 4–6 animals per cage under a 12-h light–dark cycle with ad libitum access to food and water. Survival surgeries were carried in aseptic conditions and under isoflurane anesthesia (4% induction, 2% maintenance; Med-Vet International, Mettawa IL, USA) and postoperative pain was managed with buprenorphine (s.c., 0.05 mg/kg; McKeson Medical Surgical, Irving TX USA). Terminal procedures were carried under an overdose of Euthasol (i.p., >100 mg/kg; Med-Vet International, Mettawa IL, USA). All euthanasia procedures followed the guidelines from the American Veterinary Medical Association (AVMA). We also followed Animal Research: Reporting of In Vivo Experiments (ARRIVE) guidelines.

### Animals

Wild type (WT) mice (C57Blk/6J) of both sexes were obtained from The Jackson Laboratory (Bar harbor, ME USA). Both sexes were used, but we did not detect any differences or trends and therefore both sexes were pooled together. All the animals were 8 to 10-weeks old at the start of the experiment. In the experiments described we used 96 animals. Another 19 animals were used for toxin titration until we obtained a mild dose that tetanized only one muscle for a brief period of time, without causing limb paralysis or systemic effects. The animals used for titrating the toxin were euthanized by carbon dioxide inhalation following AVMA approved methods. These animals showed different levels of tetanus, and we applied standard humane endpoints: loss of >25% of body weight, decreased body condition, leg inflammation, leg pain, lethargy, labored breathing, hunched posture, piloerection, dehydration, jaundice, cyanosis, and anorexia.

### Experimental Protocols

This study includes results from two different multilayered animal experiments. *Experiment 1 (Figure 1): Muscle reinnervation after blocking synapses with Tetanus toxin (TeTx).* In this experiment the animals underwent a muscle injection in the left tibialis anterior (TA) muscle of either TeTx or vehicle (bacteriostatic sterile saline, Med-Vet International, Mettawa IL, USA). TeTx (Lot#19050A1) was obtained from List Biologicals (Campbell, CA, USA). Seven days post-injection a sciatic nerve crush or sham surgery was performed at mid-thigh level, except for mice used for assessing TeTx effects in the absence of nerve injury. Experimental animals were split into four groups: a sham group (saline and non-crushed, n=18), a crush group (saline and crushed, n=16), a tetanus group (TeTx and non-crushed, n=17), and a tetanus-crush group (TeTx and crushed, n=26). All animals were tested pre-and post-TA TeTx-injections and nerve surgeries to assess EMG activity in the TA and lateral gastrocnemius (LG) in response to pressure applied to the foot using a calibrated custom-made pressure (PP) probe. This test allowed identification of positive tetanization in TeTx injected animals by comparing EMG activity before and after TeTx injection. Tetanization levels in individual animals were not uncovered until the end of the study and therefore the experimenter performed all tests and analyses blinded to the toxin action. Exclusion and inclusion sets and criteria are fully explained in Figure 1. Most animals underwent testing of basal muscle (M)-responses at the date of the nerve injury and prior to nerve crush or sham surgery (when appropriate). Functional muscle reinnervation was analyzed 10-, 20- or 30-days after nerve injury by monitoring recovery of M responses to sciatic nerve stimulation. A group of animals at each time post-injury was euthanized to harvest muscles and analyze muscle reinnervation by motor axons histologically. For anatomical re-innervation we analyzed only muscles at 30 days post-injury.

**Figure 1:**
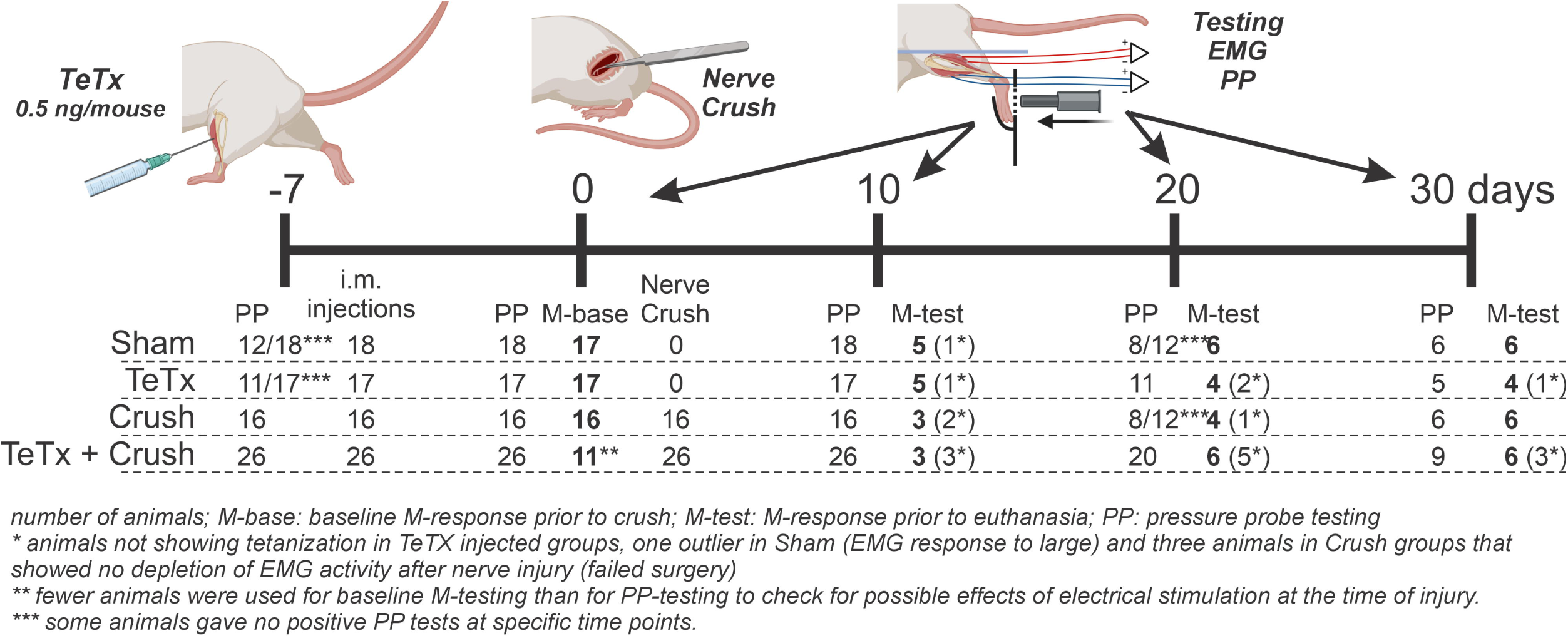
Experiment timeline and experimental groups. Four experimental groups were analyzed (Sham nerve surgery; TeTx: TA injected with TeTx but no sciatic nerve surgery; Crush: sciatic nerve crush but not injected with TeTx; TeTx + Crush: TA injected with TeTx in the TA and the ipsilateral sciatic nerve crushed). TeTx injections were performed 7 days prior (day -7) to sciatic nerve surgeries (day 0). Pressure probe tests were performed before the injection of toxin at day -7, just before nerve crush at day 0 and thereafter at 10, 20 and 30 days after nerve surgery. EMG muscle responses (M- tests) were performed just before the injection and thereafter at each survival date. Numbers indicate the number of animals prepared and tested. Animals discarded from the final analyses area indicated with asterisks. Each asterisk explains the reasons for removal from final analyses. Numbers in bold indicate the final N at each time point for the analyses in Figure 2. The animals undergoing M-tests at 10, 20 and 30 survival times were euthanized thereafter. *TeTx: tetanus toxin; TA: Tibialis Anterior; EMG: Electromyography; M-test: Muscle test responses to electrical stimulation of the nerve*.

*Experiment 2: Effect of TeTx on inhibitory and excitatory synapses on TA motoneurons*. In this experiment the animals underwent bilateral intramuscular injections of TeTx and Fast Blue in the left TA and saline and Fast Blue in the right TA. Fast Blue (Polysciences Inc, Warrington, PA USA) was injected at 1% in saline (2-4 µl). These animals also underwent bilateral PP testing prior to TeTx injections and then 7 days after injection to confirm tetanization (Supplemental Figure 1). Following PP testing the animals were euthanized as above. We injected two litters (#632 and #747) for a total of 17 animals. In one litter we observed animals with either none or effective tetanization, while in the second litter animals were either no or mildly tetanized. Although this was serendipitous it allowed us to confirm synaptic changes in controls (contralateral) with two different effects of TeTx. This pseudo dose-response analysis further strengthened conclusions. To avoid interpretation challenges due to the use of the contralateral side, two further animals were injected with saline and Fast Blue in the left TA and analyzed 7 days later. These animals did not undergo PP testing. We selected the animals with optimal Fast Blue labeling resulting in 5 controls (2 non-injected, 3 injected with TeTx contralaterally), 3 animals showing clear tetanization, 3 animals showing mild tetanization (Supplemental Figure 1). All PP testing was done in lightly anesthetized animals (0.25% isofluorane). All M-response testing, muscle injections and nerve surgeries were done in the deeply anesthetized animals (isofluorane: 4% induction, 2% maintenance). Euthanasia was performed prior to transcranial perfusion with fixatives and obtained by an overdose of Euthasol (>100 mg/kg).

### Tetanus Neurotoxin Injections

Mice were deeply anesthetized with isofluorane (4% induction, 2% maintenance in O_2_) and prepared for aseptic surgery. The left tibialis anterior (TA) muscle was exposed through a small skin incision and injected with 0.5 ng of TeTx or saline delivered in 1-2 µl. Some animals were co-injected with 1% Fast Blue in the same session. The skin was closed either with a drop of Vectabond (Santa Cruz Biotechnology, Dallas, TX USA) or by one or two 5-0 sutures and allowed to recover from anesthesia prior to returning to the home cage. Prior to surgery, all animals were given 0.05 mg/kg buprenorphine (subcutaneous) to manage post-operative pain.

### Sciatic Nerve Crush or Sham Surgeries

All nerve surgeries were performed under isoflurane anesthesia (4% induction, 2% maintenance in O_2_) using aseptic surgery conditions and the animals were given preoperative 0.05 mg/kg buprenorphine (subcutaneous) to manage post-operative pain. The left sciatic nerve was exposed by a mid-thigh skin incision and muscle separation of the overlying biceps femoris. After gently isolating the nerve from the surrounding tissue it was crushed with a pair of fine #5 forceps for 5 seconds. Nerve crush was verified by a translucent section of the nerve, approximately half-way between the sciatic notch and the trifurcation of the sciatic nerve. The wound was then closed by suturing the biceps (if necessary) with 5-0 absorbable sutures and the skin with non-absorbable ones. The animals were then removed from anesthesia. Sham animals had the sciatic nerve exposed, but not crushed.

### Pressure-probe stimulation

The extent of the tetanization was investigated prior to muscle injections, nerve injuries, and 10-, 20- and 30-days post-injury. Mice were anesthetized with isoflurane (as above) and then placed into a sling (RF S1, Lomir Biomedical Inc.) with leg holes to allow for unrestrained movement of the hind limbs. The foot was then stabilized in a holder which allowed the mouse to still flex its leg muscles while also allowing for the probe to apply pressure to the bottom of the foot, see Figure 2A for schematic. Once the mouse was restrained in the device, fine wire electrodes were inserted into the TA and LG muscles via 25-gage needles for recording M-responses and grounding wire was inserted in the back of the animal under the skin, as described below. After electrode insertion, isoflurane was reduced to 0.25% to sedation with minimal anesthesia. Mice would wake up around 2 minutes after reducing the isoflurane to this level and as such, testing was performed before this recovery would occur. Pressure was applied at 10 g, 20 g, and 30 g of force to the center of the foot and the muscle response was recorded. Recordings were obtained by plugging the free ends of EMG wires into a pre-amplifier that fed into custom-made optically isolated differential amplifiers at a gain of x1000 and sampled at 1 kHz. The signals were digitized with LabView software (National Instruments, Austin, TX USA). Each pressure level was applied 5 times with 5 seconds of recovery in between applications. LabView software was used to run our pressure-probe and acquired recordings. The records were filtered using a second order Butterworth bandpass filter, full wave rectified, and averaged using LabView. The average rectified voltages in a defined response window were quantified for each mouse (see Figure 2C). Response windows were determined as the time from when the probe started applying force to the foot until the end of the EMG.

**Figure 2:**
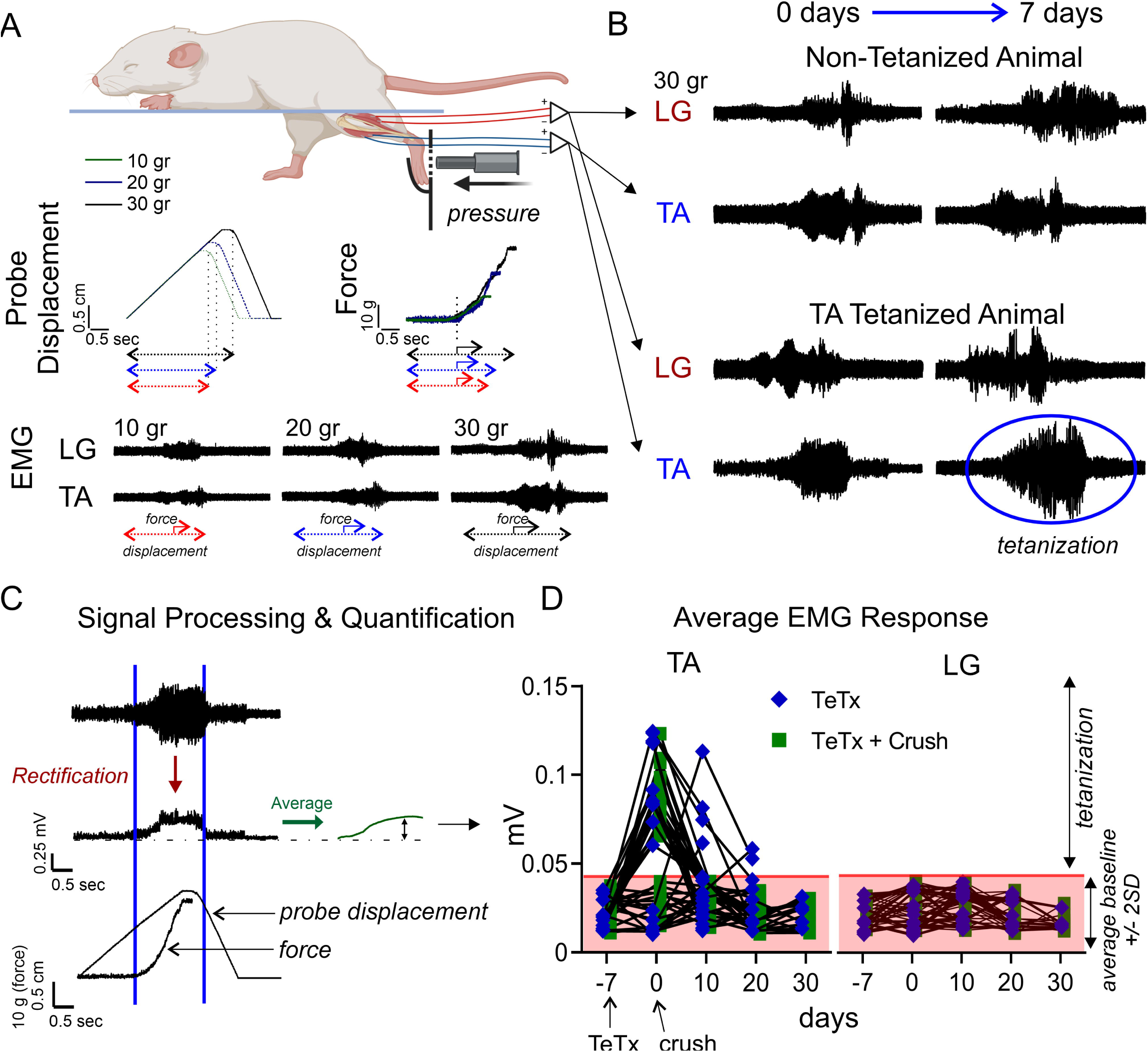
Validation of tetanus toxin effects. **A,** Experimental setup for applying calibrated pressures to the foot using a custom probe while recording simultaneously reactive foot force and TA and LG EMGs. The time course of probe displacement and force are shown with the TA and LG EMGs during foot displacement and force generation as indicated. Three different pressure forces were applied each resulting in different displacements and forces (force cannot be measured when the probe is retracted and loses contact with the foot). **B**, EMG TA and LG muscle responses before (0 days) and 7 days after injecting the left TA with saline (top records) or TeTx (lower records). Highlighted with a blue ellipsoid is the enhanced response of the TA 7 days after injection of TeTX. **C**, EMG records were rectified, integrated across time and averaged across 5 trials. The average amplitude was calculated for the EMG record from the time the force record starts to the time the EMG record ends (shortly after the probe is removed from the foot, as indicated in the force record). **D,** Average PP EMG responses in single animals across the experimental timeline (Figure 1). Only animals injected with TeTX are shown. An EMG amplitude window (grey area) was made with the average response +/- 2 standard deviations of sham animals (no nerve injury, no TeTX). Responses above this threshold were considered positive tetanization. Note that all animals in TeTx + Crush group drop to almost 0 after the nerve injury. Some animals in the TeTX group show delayed tetanus. Individual animals’ data lines stop when the animal is euthanized at scheduled survival time points. The ipsilateral LG shows no tetanization. *TeTx: tetanus toxin; TA: Tibialis Anterior; LG: Lateral Gastrocnemius Muscle; EMG: Electromyography; PP: Pressure-Probe*.

### M-response recording

The extent of functional muscle reinnervation was investigated at 10-, 20- or 30-days post-injury by recording M responses after stimulation of the sciatic nerve proximal to the injury. This testing was performed under isoflurane anesthesia (4% induction, 2% maintenance). The sciatic nerve was exposed at mid-thigh as above and bi-polar stimulating electrodes placed on the nerve proximal to the original injury. A Teflon coated stainless steel 0.003 mm diameter insulated wire was de-insulated for 5 mm at the end of the wire and inserted under the skin along the back for grounding. Fine wire electromyography (EMG) electrodes, in which the insulation was removed from the distal 1 mm of the recording tips, were inserted into the TA and LG muscles via 25-gage needles. The free ends of each wire were de-insulated and inserted into a pre-amplifier that fed into our custom-made differential amplifier. All signals were amplified (x1000), bandpass filtered (30-1000 Hz), digitized and collected with LabView software. Electrically evoked EMG activity was recorded in response to sciatic nerve stimulation (0.1 ms electrical pulses). EMG recordings were sampled at 1 kHz. Short latency direct muscle (M) responses (or compound muscle action potentials, CMAPs) are produced by motor axon activation during nerve stimulation. Stimulus intensity (ranging from 0.25 V to 5 V) was incrementally increased until a maximal amplitude direct M response was recorded. The maximal M response was used to represent the maximum extent of functional motor reinnervation of the muscle. Using custom LabView software recordings were filtered using a second order Butterworth bandpass filter, full wave rectified, and averaged. The average rectified voltages in a defined M response time window were quantified for each mouse (see Figure 3C). The M responses were recorded prior to sciatic nerve crush injury and again at the mouse endpoint of either 10-, 20- or 30-days post-injury.

**Figure 3:**
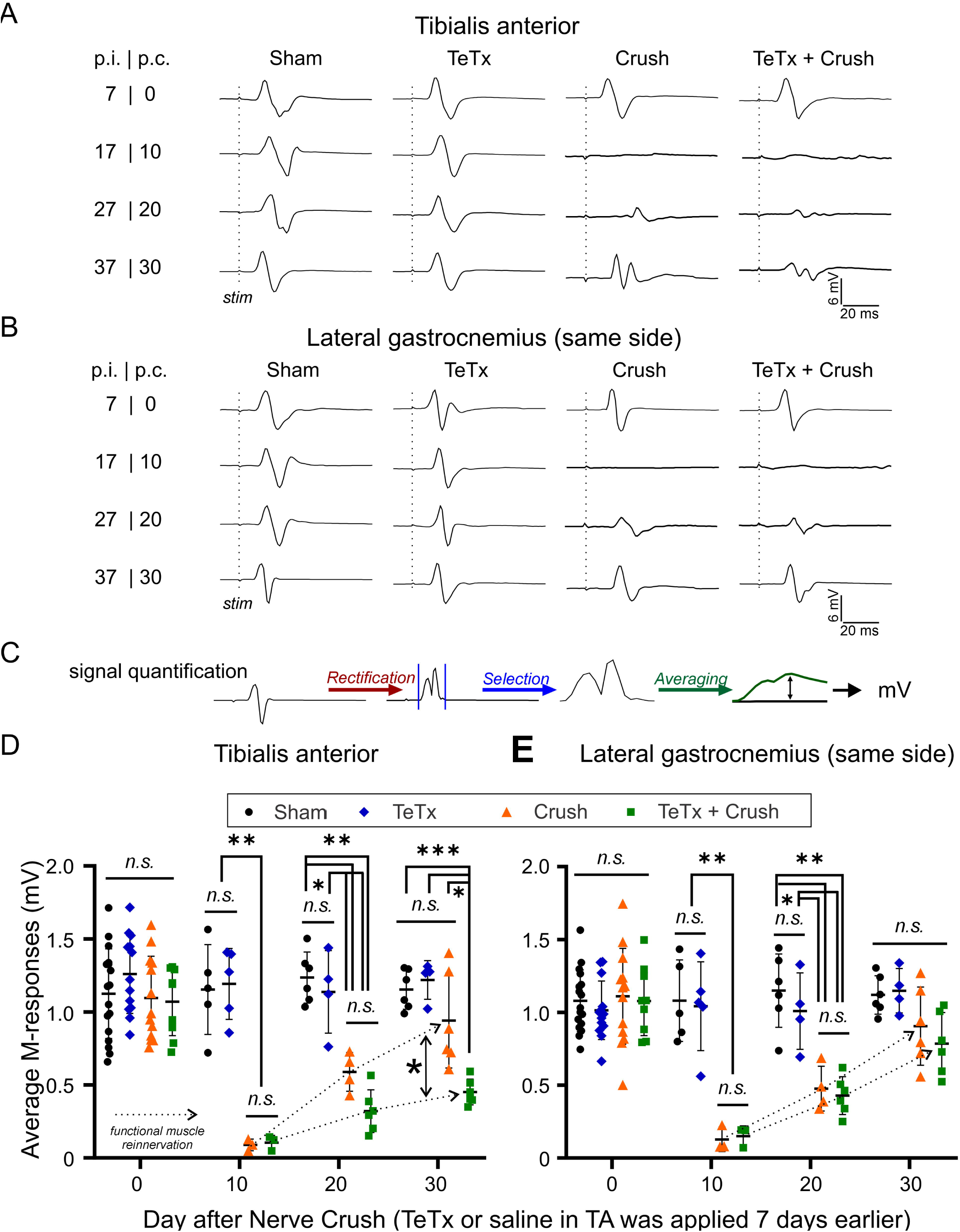
Muscle reinnervation with and without TeTx treatments estimated by recovery of M-responses. **A,** EMG records from TA in the four experimental groups at different time points. p.i.: days post injection of TeTx; p.c.: days after sciatic nerve crush surgery. Stimulation times are indicated with dashed lines. Records in Sham and TeTx animals show no change in M-responses (no effects of toxin at the neuromuscular junction). M-responses disappear after nerve crush and then partially recover by 30 days after nerve injury. Animals injected with TeTx prior to nerve crush show delayed recovery of M-responses. In all cases we show maximal M-responses. **B**, Same records for the ipsilateral LG. Recovery after crush is the same whether the ipsilateral TA was injected with TeTx or not. There are no bystander effects. **C**, Process of signal manipulation to average M-response across trials (see Methods for explanation). **D, E,** Average M-responses at different experimental time points in each of the four experimental groups in TA (**D**) and LG (**E**). Each data point represents one animal. The N is explained in Figure 1 with the experimental design and timeline. Average and standard deviation (error bars) are indicated. In both TA and LG, M-responses drop to almost 0 at 10 days post-injury and then start to recover. Recovery is significantly slower in the injected TA compared to non-injected TA and to LG muscles ipsilateral to TA muscles injected or not injected with TeTx. Data was compared using a two-way ANOVA for time after injury and experimental group. The results showed significant differences (p<0.0001) for both variables and their interactions. All statistics are fully reported in supplementary Tables 2 (for D) and 3 (for E). Asterisks denote the result of Tukey’s pair-comparisons performed among experimental groups at each survival time: *p<0.05, **p<0.01; ***p<0.001. There is a significant delay in recovery of the M-reposes in the TeTx injected TA. The dotted arrows denote the functional muscle reinnervation time-course in each muscle and experimental group. *TeTx: tetanus toxin; TA: Tibialis Anterior; LG: Lateral Gastrocnemius Muscle; EMG: Electromyography; PP: Pressure-Probe; M-response: Muscle responses to electrical stimulation of the sciatic nerve*.

### Tissue collection, processing, and immunocytochemistry

Following functional assessment, the mice were overdosed with Euthasol (>100 mg/kg) and transcardially perfused with heparin-saline followed by paraformaldehyde (PFA; 4% in 0.1M phosphate buffer). The spinal cord and the TA and LG muscles were harvested and postfixed overnight in 4% PFA before cryoprotection in 30% sucrose for at least 24 h. The muscles were initially compressed between two microscope slides in 0.01M Phosphate saline with 0.9% NaCl (PBS) (to prevent the muscles from drying out) under approximately 5 pounds of weight for 2-3 hours. Following compression, the muscles were embedded in OCT and frozen. Cryostat sections were cut at 60-µm thickness and collected onto glass slides. After washing 5 times in 0.01 M PBS, the sections were incubated for 2 h in 0.02 M sodium citrate with 20% triethanolamine, 0.5% SDS, and 0.5% Tween-20 (pH 6, room temperature) for antigen retrieval. The slides were washed again (3X in 0.01 M PBS) and then blocked for 1 h in 5% Normal Donkey Serum, diluted in PBS with 0.3% Triton (PBST). Sections were incubated overnight at room temperature with α-bungarotoxin (α-BTX) conjugated to Alexa Fluor 647 (Thermo Fisher Scientific, Waltham, MA USA) and primary antibodies against neurofilament-H (NF-H) to label the axonal cytoskeleton in thick motor axons and against vesicular acetylcholine transporter (VAChT) to label the presynaptic motor nerve terminals (Table 1). VAChT-IR was revealed with donkey Cy3-conjugated species-specific anti-goat IgG secondary antibodies and NF-H with donkey FITC-conjugated species-specific anti-chicken IgY secondary antibodies (dilution 1:200; Jackson Immunoresearch, West Grove, PA USA). After washes in PBS the sections were mounted in Vectashield (Vector Labs, Newark, CA USA) and imaged in Olympus FV1000 confocal microscope. We obtained Z-stacks through the 60 µm thick sections to obtain 50 motor endplates per sample for quantification.

**Table 1.** Primary antibodies used for immunohistochemistry.

| Antigen | Immunogen | Host/type<br>Dilution | Manufacturer | Catalogue #<br>Lot# | RRID# |
| --- | --- | --- | --- | --- | --- |
| VGLUT1 | Recombinant protein corresponding to residues near the carboxy terminus of rat VGLUT 1 | Guinea Pig / Polyclonal 1:1000 | Synaptic Systems | 135-304<br>Lot# 1 53 | AB_887878 |
| VGLUT2 | Synthetic peptide corresponding to residues near the carboxy terminus of rat VGLUT2 | Guinea Pig / Polyclonal 1:1000 | Synaptic Systems | 135-404<br>Lot# 3 39 | AB_887884 |
| VGAT | Recombinant protein corresponding to residues near the amino terminus of rat VGAT | Guinea Pig / Polyclonal 1:500 | Synaptic Systems | 131-004<br>Lot# 5 58 | AB_887873 |
| GAD65 | GAD partially purified from chicken brains | Mouse / Monoclonal GAD-6 clone 1:10 | Developmental hybridoma bank | GAD-6<br>Lot# 2/1/2020 | AB_528264 |
| VAMP1 | Synthetic peptide corresponding to AA 2 to 14 from rat Synaptobrevin1 | Rabbit / Polyclonal 1:500 | Synaptic Systems | 104-002<br>Lot# 1 16 | AB_887807 |
| VAMP2 | Synthetic peptide corresponding to residues near the amino terminus of rat Synaptobrevin2 | Rabbit / Polyclonal 1:500 | Synaptic Systems | 104-202<br>Lot# 1 16 | AB_887810 |
| NeuN | Purified cell nuclei from mouse brain | Mouse / Monoclonal 1:250 | EMD Millipore | MAB377<br>Lot#3278580 | AB_2298772 |
| NF-H | Native NF-H purified from bovine spinal cord | Chicken / Polyclonal 1:1000 | EnCor Technologies | CPCA-NF-H<br>Lot#7575-8 | AB_2149761 |
| NF-H | Native NF-H purified from bovine spinal cord | Chicken / Polyclonal 1:1000 | Aves | NFH<br>Lot# 7857983 | AB_2313552 |
| VACHT | A synthetic peptide from human VACHT conjugated to an immunogenic carrier protein | Goat / Polyclonal 1:5000 | Invitrogen (Thermo Fisher Scientific) | OSG00003W<br>Lot# XA3481392 | AB_2068697 |

The spinal cords were sectioned in a transverse plane on a freezing sliding microtome at 50-µm thickness and collected free-floating. The sections were washed three times in 0.01 M PBS with 0.3% Triton (PBST) and blocked in normal donkey serum (10% in PBS with 1% Triton) for 1 h. The tissue was then incubated overnight, while shaking at room temperature, with a mixture of appropriate primary antibodies diluted in PBST. The antibodies used, their sources, and RRID numbers are listed in Table 1. Sections were reacted with primary antibodies against either an excitatory (VGLUT1 or VGLUT2) or inhibitory marker (VGAT) and either VAMP1 or VAMP2 and NeuN. Sections were then washed in PBST and incubated with appropriate fluorescent-coupled species-specific anti-IgG secondary antibodies (1:100 in PBST, Jackson Laboratories, West Grove, PA USA). Immunoreactivity (IR) to VGLUT1, VGLUT2 or VGAT was visualized after incubating with FITC-coupled antibodies, VAMP1-IR and VAMP2-IR with Cy3-coupled antibodies, and NeuN-IR with Cy5-coupled secondary antibodies. TA motoneurons were retrogradely labeled with FB. In a few sections we combined VAMP2, GAD65 and VGLUT1 antibody to confirm VAMP2 expression inside GAD65 terminals surrounding VGLUT1 synapses (i.e., presynaptic axons). After washing, the sections were coverslipped with Vectashield (Vector Labs) and imaged (four channels) on an Olympus FV1000 confocal microscope. Image Z-stacks were obtained with 20X (NA, 0.75) objectives and then at high-magnification using a 60X objective with 2X digital zoom (NA, 1.35, oil-immersion) for quantification of VAMP1 and 2 content inside different types of neurochemically identified excitatory and inhibitory synapses.

### Automated quantification of NMJ re-innervation

α-BTX-labeled NMJs were analyzed using an automatic procedure developed in Image J (Figure 4B). First the image stack was transformed into the average intensity projection and using α-BTX fluorescence to identify the motor endplates, a rectangular ROI was drawn around each NMJ that was in phase with the horizontal plane of the image (out of phase or NMJs perpendicular to the image were not used). Using the drawn rectangle, the mean background intensity plus one standard deviation was subtracted from the NMJ for each fluorescent channel. After background subtraction, the resulting α-BTX fluorescence was used to trace the NMJ. The overlap of VAChT fluorescence and α-BTX fluorescence in the trace was then used to determine the reinnervation of the motor neuron. Fully innervated NMJs had >50% coverage, partially innervated NJs displayed >10% but less than 50% coverage. We analyzed 50 NMJs in the TA and LG of the left leg in each animal in each of the four experimental groups described in Experiment 1 (above). For this study we selected the 30 days survival time point: n=6 Sham; n=5 TeTx; n=6 crush; n=6 TeTx+Crush.

**Figure 4.**
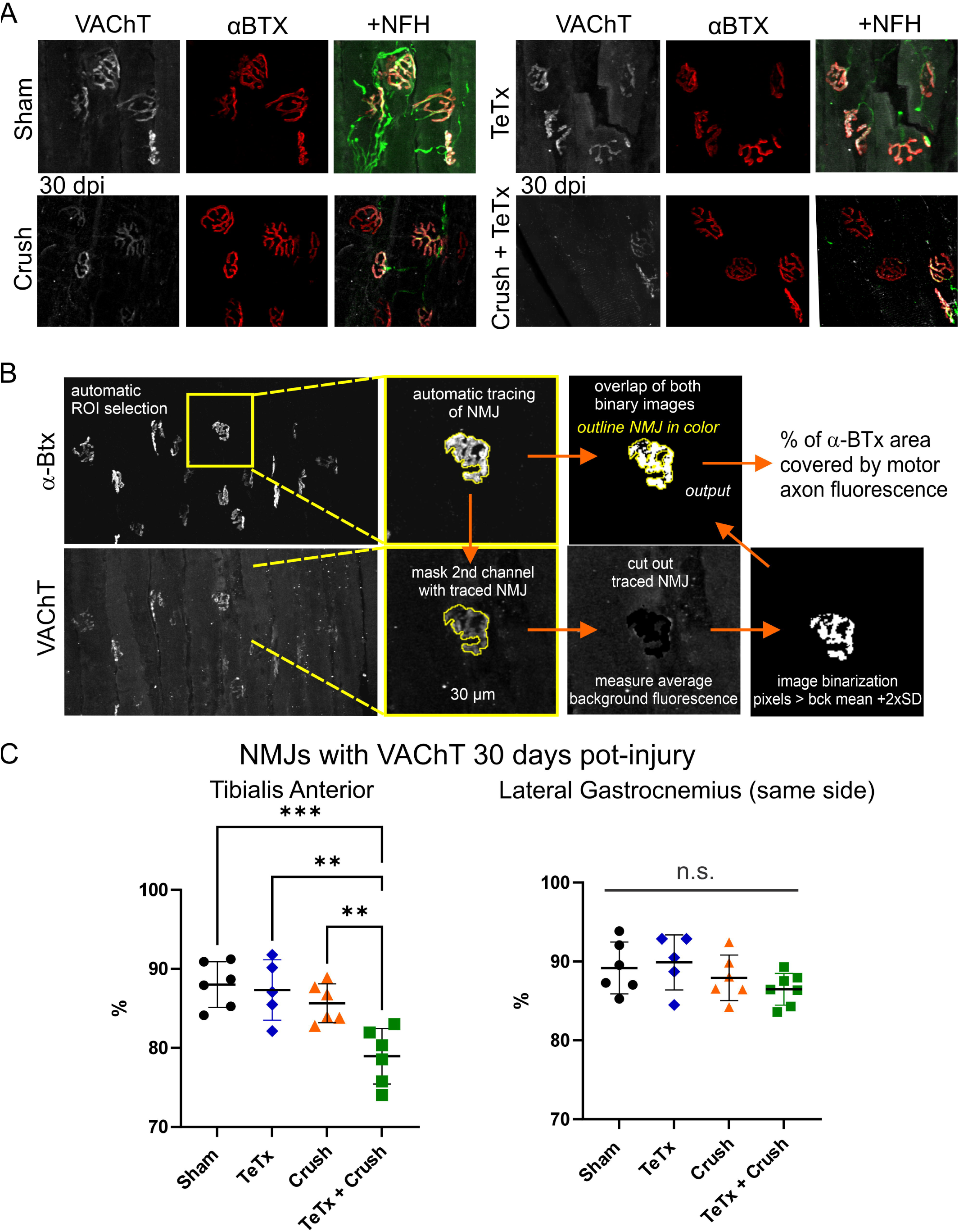
Recovery of anatomical reinnervation of neuromuscular junctions (NMJs) by motor endplates 30 days after nerve crush in the TA. **A,** Staining of motor endplates with antibodies against vesicular acetylcholine transporter (VAChT, white, Cy5), the postsynaptic NMJ receptor fields by binding of α-bungarotoxin coupled to Alexa-555 (αBTX, red) and the motor axons with antibodies against the phosphorylated heavy neurofilament chain (NFH, green, FITC). All three stains are shown for all four experimental groups 30 days post injury (dpi) in animals with sciatic nerve crush and at the same time point for sham and TeTx no-injury animals. **B,** Method for quantification of αBTX NMJ coverage by VAChT positive motor endplates (see Methods for full explanation). **C,** Percentage of NMJs covered with VAChT in the injected TA (graph at the left) and ipsilateral LG (graph at the right). Each dot represents data from one animal across 50 NMJs. Average and standard deviations (error bars) are indicated. In the TA there are significantly fewer innervated NMJs in animals injected with TeTx compared to all other animals, including animals recovering from nerve crush. No significant differences were found in the ipsilateral LG. Statistical comparisons were performed using one-way ANOVAs across experimental groups (p=0.0004 for TA; p=0.2171 for LG) followed by post-hoc pairwise comparisons using Tukey tests: **p<0.01, ***p<0.001. All details on stats are fully reported in supplementary table 3. *VAChT; Vesicular Acetylcholine Transporter, NMJ: Neuromuscular Junction; NFH: Neurofilament Heavy subunit; TeTx: Tetanus Toxin; αBTX: α-bungarotoxin; Cy5: Cyanine5; FITC, Fluorescein isothiocyanate; TA: Tibialis Anterior; LG: Lateral Gastrocnemius Muscle*.

### Quantification of VAMP content inside VGLUT1, VGLUT2 and VGAT synapses

Following image acquisition of the motor neurons, colocalization of VAMPs with the different synaptic markers were analyzed using Image J. Using the ‘wand tracing tool’ every structure marked for VGLUT1, VGLUT2 and VGAT (FITC in green) was thresholded, and their area and length measured, after which the bouton was outlined, and the average mean grey value of the VAMP1 or VAMP2 fluorescent signal (Cy3 in red) was measured within the outline. The synapses analyzed were all contacting the cell bodies of neurons in confocal planes with a nucleolus present. In total 268, 1,564 and 1,201 boutons in contact with TA motoneuron cell bodies were analyzed for VGLUT1, VGLUT2 and VGAT synapses (Supplementary Table 10). This represents an average of 5.7 ±0.2 (±SD) VGLUT1, 15.5 ±1.2 VGLUT2 and 16.4 ±5.7 VGAT boutons per motoneuron single confocal plane cross section. In this study we used only control and TeTx animals (no nerve injured animals). The TeTx animals were divided into animals that displayed positive or weak/negative tetanization (Supplementary Table 10 and Supplemental Figure 1). Boutons were then analyzed for the presence or absence of VAMP1 or VAMP2 to determine changes due to the injection of TeTx in the muscle. Those synaptic boutons where VAMP1 or VAMP2 signal had a higher intensity than the average ROIs signal +2 standard deviations were considered positive for their corresponding VAMP and a percentage was obtained. To estimate the average density of VAMP1 or VAMP2 we included in the analyses only synapses that were considered positive.

### Antibody properties and specificity

All primary antibodies used are listed in Table 1 with antigen, species, purification and RRID numbers described. The specificity of vesicular neurotransmitter transporters has been extensively tested by our group including testing in knockout tissues (KOs), comparison of different antibodies against the same antigen raised in different species and co-localization with genetic labeling of neurons expressing the same proteins (Siembab *et al*., 2010; Alvarez *et al*., 2011; Siembab *et al*., 2016). The specificity of the VAMP1 antibody used was tested in KO tissues (Vuong *et al*., 2018) and similarly VAMP2 (Wang *et al*., 2022). NeuN antibodies correspond to the original A60 clone characterized by Mullen and colleagues (Mullen *et al*., 1992) an later identified as Fox-3 by biochemical/genetic means (Kim *et al*., 2009). GAD65 antibodies correspond to the GAD-6 clone fully characterized by Chang and Gottlieb using western blots (Chang & Gottlieb, 1988). GAD-6 labeling in the ventral horn of P-boutons on VGLUT1 synapses replicates genetic labeling of these same boutons containing GAD65 (Mende *et al*., 2016). Two different antibodies were used against neurofilament phosphorylated heavy protein resulting in similar immunostains of large peripheral axons.

### Microscopy and Image handling

All images were obtained in an Olympus Fluoview 1000 laser scanning confocal microscope. All images were obtained at 1024 x 1024 image pixel size using a 10X (NA:0.40), 20X (NA:0.75) and 60X (NA:1.35, oil) objectives corresponding to respectively, 1.242, 0.621 and 0.207 µm/pixel. The number of z-planes displayed in each individual image is reported in figure legends. Immunoreactivity was usually captured throughout the whole tissue thickness (50 or 60 µm). The z-steps used were 1.5, 1, and 0.5 µm respectively for the 10X, 20X and 60X objectives. The total number of optical planes captured depended on tissue shrinkage and surface unevenness. Figures were composed in Corel Draw (ver. 16 to 24). Image brightness, gamma and contrast were optimize using either Image Pro Plus (Ver 7, Media Cybernetics, Rockville, MD, USA) or Image J. All manipulations were minimal, applied to the whole image and did not alter information content.

### Statistics

When data distributed normally (as determined Shapiro-Wilk test) we used two-way ANOVAs to compare time after injury and experimental group. Multiple pair-wise comparisons following significance in ANOVA tests were compared using post-hoc Tukey’s tests. When data were not normally distributed (for example when analyzing VAMP content) we used Kruskal-Wallis multiple comparisons ANOVA followed by Dunn’s test for pair wise comparison. Effect sizes in pair comparisons was estimated using Hedges’ g and the corrected formula for small samples.

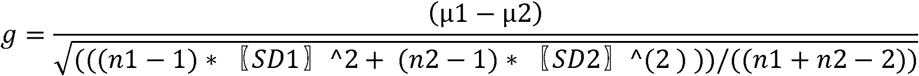

Significance was set to p < 0.05. All statistical comparisons were made using GraphPad Prism (version 9.5.1, GraphPad Software, Boston, MA, USA). All statistical parameters and results are reported in supplementary data tables 1 to 10. When necessary, power was tested in SigmaPlot 12.0 (Systat Software, Inc. Palo Alto, CA). Power goal was 0.8 for α = 0.05.

## Results

### Low doses of tetanus toxin injected in one muscle (tibialis anterior) induces motor pool-specific tetanization with minimal global effects

Tetanus toxin (TeTx) effects are dose-dependent and range from generalized tetanus characterized by muscle spasms, whole limb rigidity, autonomic dysfunction and eventually animal death, to reversible focal spasticity localized to single muscles (Goonetilleke & Harris, 2004; Kutschenko *et al*., 2012; Megighian *et al*., 2021). This study sought a localized and short-acting tetanization of just one motor pool. TeTx doses and lots were thoroughly tested to avoid generalized tetanus, limb spastic paralysis or inflammation. The lot of TeTx used in this study was obtained from List Biologicals (Lot#19050A1) and had a reported endotoxin content of 10 EU/mg, with full VAMP cleavage detected within 3 hours using 200 mM of TeTx and 20 µM VAMPtide and GT1b ganglioside binding reported with 6.5 µg/µl in hemaagglutination assay (per manufacturer data). We injected TeTx in hindlimb muscles of 2- to 3-month-old C57/BL6 mice at doses from 0.005 to 5 ng/Kg. We discarded doses that resulted in animal death or evolved into long-lasting paralysis of the injected limb or inflammation, sometimes requiring euthanasia. We selected a low dose that showed no behavioral deficits during normal ambulation in the home cage: 1 µl (two 0.5 µl injections) of 0.5 ng/µl TeTx (diluted in sterile saline) for a final dose between 0.03 and 0.02 ng/Kg depending on body weight (16 to 25 grams) injected unilaterally (left) into the tibialis anterior (TA) muscle.

For analyses of nerve regeneration we divided the mice in four experimental groups: sham nerve surgery animals with no TeTx (Sham, n=18), TeTx animals with sham surgery (TeTx, n=17), animals with a sciatic nerve crush, but no TeTx (Crush, n=16) and animals that underwent a sciatic nerve crush and also the TA was injected with TeTx (TeTx + Crush, n=26) (Figure 1). The level of tetanization of the TA was tested using a custom-made pressure probe (PP) to deliver a calibrated external force to the foot pad while animals are partially restrained in a full body sling lifting the legs from the ground such that there are no weight forces on limb joints. This stimulus evoked a stabilization response at the ankle characterized by co-contraction of ankle flexors (TA) and extensors (LG) (Figure 2A). During PP tests we recorded simultaneous electromyography (EMG) from TA and LG while applying 3 different forces (10g, 20g, 30g) at a speed of 3 mm/s for a maximum displacement of 15 mm. EMG responses to the probe build slowly and then stop a few milliseconds after the probe is removed. Signal-to-noise ratio was highest in 30g trials, and this force was chosen for subsequent analyses (Figure 2A). Motor pool disinhibition was identified as an increased EMG response. This was detected at 7 days after TeTx, specifically in injected TA muscles (Figure 2B). To quantify the EMG response, we analyze the average amplitude of the rectified signal (5 consecutive responses recorded with 5s delay between tests) throughout a time window initiated by the first detected change in force and ending with the EMG burst (force could not be measured once the probe separated from the foot at the start of the probe retraction) (Figure 2C). We calculated a baseline response from PP tests in all animals 7 days before the nerve crush or sham surgery and before injecting TeTx or saline (vehicle) in the muscle (day -7 in Figures 1 and 2). Animals were retested the day of nerve surgery (that is 7 days after TeTx injection) and thereafter every ten days. TeTx-injected animals showing responses above the average maximal baseline amplitude +2 standard deviations were considered tetanized (Figure 2D). Fifty-six percent of animals injected with this low dose of TeTx in TA showed increased EMG responses in the TA muscle 7 days post injection (n=24 out of 43). None showed increased EMGs in the LG muscle of the same leg, confirming that TeTx effects were localized to one muscle and motor pool. Sixty percent of animals showing tetanization returned to baseline 17 days after toxin injection (corresponding to 10 days after nerve injury), while the rest had returned to baseline when tested at day 27 post-toxin injection (i.e., 20 days after the nerve injury). This suggests toxin effects were present at the time of the injury and during the first week of regeneration, with effects lasting more than one week in a few animals. Forty-four percent of injected mice showed no tetanization 7 days after toxin injection, but 3 animals (18%) in the TeTx injected cohort with no nerve crush showed delayed responses at day 17. Late responders in the TeTx cohort (without nerve injury) were included in final analyses. All testing was performed blind, and the identity of mice in different experimental groups revealed only in final analyses. Late responders were obviously not detected in TeTx + Crush group because muscle disconnection abolished any EMG responses. Moreover, at this timepoint KCC2 is downregulated in axotomized motoneurons likely abolishing the inhibitory actions of GABA and glycine. Animals in the TeTx + Crush group showing no TeTx effect before nerve injury were removed from final analyses (these animals are indicated in Figure 1 with one asterisk). Late or lack of tetanized responses in some animals is expected because the very low doses used in this study. Finally, it is important to note that the extent of tetanization at 7 days varied, with some mice displaying higher or longer tetanized responses than others (Figure 2D).

In summary, we used rigorous protocols to validate specific and short-lived tetanization of just one motor pool, the TA, to compare regeneration with other pools in the same animals that were also injured after the sciatic nerve crush but did not undergo TeTx block of neurotransmitter release.

### Blocking GABA/Glycine release with tetanus toxin delays functional muscle reinnervation

To functionally investigate the time course of muscle reinnervation after injury, we analyzed muscle response EMGs (compound muscle action potentials; M-responses) after sciatic nerve stimulation in both TA (the tetanized muscle) and the LG (non-tetanized muscle in the same leg). The day of nerve surgery a baseline M response was obtained prior to the surgery. Maximal M responses were estimated by increasing voltage stimulation steps applied to the sciatic nerve (see methods). Stimulation steps stopped when M-response amplitudes plateau for 3-4 step increases. Maximal M-responses were calculated from two consecutive trials. M-responses were recorded after PP tests at 10, 20 and 30 days after nerve crush injury or sham surgery in animals with or without a TeTx injection. Animals were euthanized after measuring M-responses and the muscles and spinal cords collected for analyses (see below). Most animals therefore underwent two M-response tests: baseline at day 0 and then at their experimental end-point (10, 20 or 30 days after nerve crush injury). A few animals did not receive baseline M-testing to rule out possible effects on regeneration of the electrical stimulation used for baseline M-tests prior to the injury. No differences were observed in recovery of M-responses between animals with or without baseline M-tests done prior to the injury. This is not too surprising since electrical stimulation protocols that effectively promote regeneration are done after injury, for 30-60 minutes and at 20 Hz (Al-Majed *et al*., 2000a; Al-Majed *et al*., 2000b). This is markedly different to the electrical stimulation protocol we used during M-response tests. Animals in which we did not detect effective tetanization of TA muscles were removed from final analyses.

Representative traces of EMG recorded M-responses are shown for the TA (Figure 3A) and the LG of the same leg (Figure 3B) at different time points after TeTx injection (post-injection: p.i.) and nerve surgery (post-crush: p.c.) for all four experimental groups: Sham, TeTx (no nerve injury), Crush and TeTx + Crush. Sham and TeTx animals showed similar M-responses in both muscles at all time points indicating that TeTx, at this dose, had no discernible effects on the neuromuscular junction. As expected, ten days after sciatic nerve crush no M-responses could be elicited in either TA or LG because muscle denervation. M-responses started to recover in non-tetanized TA and LG muscles 20 days after injury and were almost fully recovered by 30 days post-injury, however, M-response amplitude recovery was delayed in tetanized TA muscles. To compare average M-responses in TA (Figure 3D) and LG (Figure 3E) for each experimental group (different symbols) and days after nerve crush (x-axis) we used two-way ANOVAs on estimates of average amplitudes obtained from the rectified and filtered EMG trace (Figure 1C, see also Methods). These estimates showed significant differences for time after injury and experimental group in both TA and LG (two-way ANOVA p<0.0001 in all cases; statistical details in Supplemental Tables 1 and 2). We further tested significance in pair-wise comparisons using post-hoc Tukey’s tests and estimated effects sizes calculating for each pair a Hedges’ g and a percentage difference (supplemental Tables for Figures 3D and 3E). Ten days after injury all animals with the sciatic nerve crushed (Crush and TeTx + Crush groups) lacked M-responses compared to Sham and TeTx injected animals with no nerve crush in both TA (p<0.01 in all cases). Effect sizes were very large (Hedge’s g from 4.1 to 5.5) with reductions ranging between 90.9 to 92.5% of Sham or TeTx EMG values. Similar results were observed in the non-injected LG 10 days after sciatic nerve crush (p<0.01 in all cases, Hedge’s g from 3.5 to 4.7; 85.5 to 87.7% of Sham or TeTx values). At twenty days after injury some M-response recovery was observed. In the Crush group, M-responses were now only 52.4 and 48.2% reduced from Sham and TeTx values, respectively (g=4.0 and 2.5 and p=0.0008 and 0.0732). However, the M-responses of the TA in the TeTx + Crush group were still depleted by 74.1 and 71.8% of Sham and TeTx values (g=5.7 and 3.9; p < 0.0001 and p=0.0193). The average M-response in TA 20-days after nerve crush in TeTx injected animals compared to non-injected animals (Crush group) (Figure 3D) did not reach significance (p=0.0741) despite large differences (estimated difference: 45.6% and g=1.9). In comparison, M-responses in the non-injected LG recovered at the same rates in animals with or without TeTx in TA. Injured animals with or without TeTx showed similar amplitude differences in the LG compared to Sham (62.7 and 58.6%; g=3.6, 3.1; p=0.0015, 0.0035) or TeTx (no injury) animals (57.5 and 52.8%, respectively; g=3.0, 2.5; p=0.0489, 0.0629). Accordingly, no significant differences were detected in LG M-responses 20 days post-injury between Crush and TeTx + Crush groups (p=0.9540). Thirty days after injury, animals receiving a nerve crush without TeTx showed recovered M-responses in TA and LG, although with large inter-animal variability (Figure 1D, 1E). On average, TA M-responses in Crush animals showed no significant differences with respect to Sham (g=0.8; p=0.4952) or TeTx (no crush) animals at this time after injury (g=1.0; p=0.3168). In contrast, TA M-responses in animals recovering from nerve crush with a TeTx injection in the TA remained reduced compared to Sham (60.9%, g=6.1 p<0.0001) or TeTx animals with no nerve injury (63.0%, g=7.1, p=0.0008). Consequently, a very significant difference was detected 30 days post-injury between recovered M-responses in animals with the TA injected with TeTx or not injected (52% differences, g=2.1, p=0.0475). At this time point M-responses were recovered in the LG in animals in which the neighboring TA was injected with TeTx (Figure 3E). Although a lingering significant decrease with the Sham group (29.8%, g=1.9) was detected in the LG of the TeTx + Crush group (p=0.0436), no significant differences were found with TeTx or Crush groups (supplemental Table for Figure 3E).

In summary, pre-injection of the TA with TeTx caused a significant delay in muscle reinnervation by TA motoneurons that was detectable at 20- and 30-days post-injury with no significant effect on non-injected muscles of the same leg. We do not expect errors in regeneration to confound this conclusion (for example, non-tetanized LG motoneurons reinnervating the TA or tetanized TA motoneurons innervating the LG) because we have shown high reinnervation specificity to these muscles in this sciatic crush-injury model (Rotterman *et al*., 2024).

### Blocking GABA/Glycine release with tetanus toxin delays anatomical muscle reinnervation

To confirm delayed reinnervation of muscle by motor axons we analyzed the re-occupancy after injury of neuromuscular junctions (NMJs, labeled with α-bungarotoxin (α-BTX) conjugated to Alexa Fluor 647) by motor end-plates (labeled with antibodies against vesicular acetylcholine transporter, VAChT) in animals and muscles with our without TeTx (Figure 3A). We analyzed 53 to 68 NMJs per animal/muscle (average=58.9 ±4.4 S.D.) in the TA and 52 to 70 in the LG (average=61.2 ±4.8 S.D.) in the same experimental groups: Sham (n=6 animals), TeTx (n=5), Crush (n=6) and TeTx + Crush (n=7). We analyzed NMJ reinnervation 30 days after nerve injury, the time point corresponding to the largest difference found in M-responses in regenerating animals with and without TeTx.

To estimate the number of re-innervated NMJs we developed a bias-free automatic processing using ImageJ (Figure 4B). All NMJs in the field were automatically threshold and outlined first, then the VAChT signal was threshold, captured and superimposed to the NMJ outline to calculate a percentage coverage. Thresholding criteria was the same for both signals: two standard deviations above the average background, calculated independently for each confocal image plane over muscle fibers outside the NMJ. Using this method, we found that when the percentage of NMJ area covered by VAChT is more than 50%, VAChT can be visualized over every compartment of the NMJ, therefore these NMJs were considered fully covered by the motor end-plate. We also calculated NMJs with VAChT coverage of >10% and ≤50%. These NMJs were considered partially cover by VAChT motor end-plates. Adding fully and partially covered NMJs we calculated all NMJs associated to a motor end-plate. Using these criteria, 60.6% ±7.8 (±S.D.) of NMJs in the TA of Sham animals were fully covered by VAChT and 88.0% ±2.9 were considered with innervation. Similar results were obtained in the LG (61.1% ±1.8 and 89.2% ±3.3). These values are lower than expected for intact NMJs. Differences in signal-to-noise ratio using αBTX binding or VAChT antibodies might explain these results because they influence the efficiency of complete automatic capture of each of the signals: complete for αBTX-labeled NMJs but only partial for VAChT-labeled motor end-plates with lower signal-to-noise ratio. Despite these labeling difficulties, the method has the advantage of being fully automatic and free of experimenter bias. Following analyses were based on NMJs with >10% VAChT coverage to more fully represent the presence of innervation, even when the signal-to-noise ratio prevents fully detecting all VAChT immunostaining.

The results (Figure 4C) show that the percentage of NMJs with innervation is similar in Sham and TeTx animals in both TA and LG. Moreover, the NMJ innervation ratio after sciatic nerve crush is also not significantly different in TA and LG 30 days after nerve crush (Tukey post-hoc pair-wise tests following a significant one-way ANOVA: vs. Sham group p=0.5806 (TA), 0.8752 (LG); vs. TeTx group p=0.8185 (TA), 0.6842 (LG); statistical details in Supplementary Table 3). This suggests effective reinnervation at this time after injury, confirming conclusions derived from M-responses. In contrast, there were significant differences in innervation after crush in tetanized TA muscles compared to controls (vs. Sham group p=0.0005; vs. TeTx group p=0.0018) or regenerating animals (vs Crush group p=0.0085). Differences in percentage of innervated NMJs range from 10% compared to controls (Hedge’s g=2.8) to 8% compared to animals regenerating from Crush (with no toxin) (g=2.2). This parallels the delayed recovery of M-responses detected in TeTx injected animals recovering from nerve crush, however the effect sizes are smaller (8-10% difference in NMJ innervation with Hedge’s g=2.2 to 2.8 vs 52-63% difference in M-response amplitude with g=2.1 to 7.1). The different effect size suggests higher reinnervation than predicted from M-responses. This difference is best explained by delayed recovery of NMJ function with respect to anatomical reinnervation by motor endplates, as previously reported (Arbat-Plana *et al*., 2023). The LG muscle of the same leg injected with TeTx in the TA recovered NMJ innervation after nerve crush similar to the Crush group and there were no statistically significant differences in innervation ratio between Sham, TeTx, Crush and TeTx + Crush groups in the LG (one-way ANOVA p=0.2171, details in statistical table for Figure 4C). These data further support the conclusion that motor axon regeneration is specifically delayed in tetanized TA motor pools.

### Tetanus toxin cleaves vesicular associated membrane protein specifically from inhibitory synapses without isoform specificity

To investigate the effects of TeTx muscle injections on synaptic inputs over targeted motoneurons, we compared VAMP expression in different types of synapses on the cell body of retrogradely labeled TA motoneurons with and without injecting TeTx in the muscle. Fast Blue (1-2%) was injected in the TA muscle simultaneously to TeTx or vehicle in the left and right-side legs of 17 animals. Analysis of TA tetanization by PP testing 7 days after injection divided the TeTx cohort into two groups, one with high tetanization and another with low or no tetanization (Supplemental Figure 1). Optimal Fast Blue retrograde labeling was found in five animals in the control side and in three animals each with high or low effect of TeTx. These resulted in three experimental groups for histological analyses: 1) control TA (vehicle injection, n=5 animals), 2) TeTx injection with low or no TA tetanization (n=3), 3) TeTx injection with clear tetanization of the TA muscle (n=3). Synaptic boutons on the cell bodies of Fast Blue labeled TA motoneurons were divided into VGLUT1, VGLUT2 and VGAT synapses (green channel) and their VAMP content (red channel) examined. Frequently, motoneuron cell bodies were also labeled with NeuN (far red channel). In control non-injected TA motoneurons, all VGLUT1 synapses contained VAMP1 and not VAMP2 (Figure 5A), while most VGLUT2 and VGAT synapses expressed VAMP1 and/or VAMP2 (Figures 5F and H and Figures 6A and D). VAMP1 and VAMP2 co-expression in single synapses was not examined because both VAMP antibodies were raised in the same species. Nevertheless, their abundance in VGLUT2 and VGAT synapses suggests frequent co-expression. VGLUT1 synapses were surrounded by small VAMP2 boutons resembling presynaptic P-boutons. Therefore, in some sections we modified our primary antibody combination to include GAD65 (green channel) with VAMP2 (red channel) and VGLUT1 (far-red channel). Every GAD65 bouton opposed to VGLUT1 terminals contained VAMP2, but never VAMP1 (Figure 5C).

**Figure 5.**
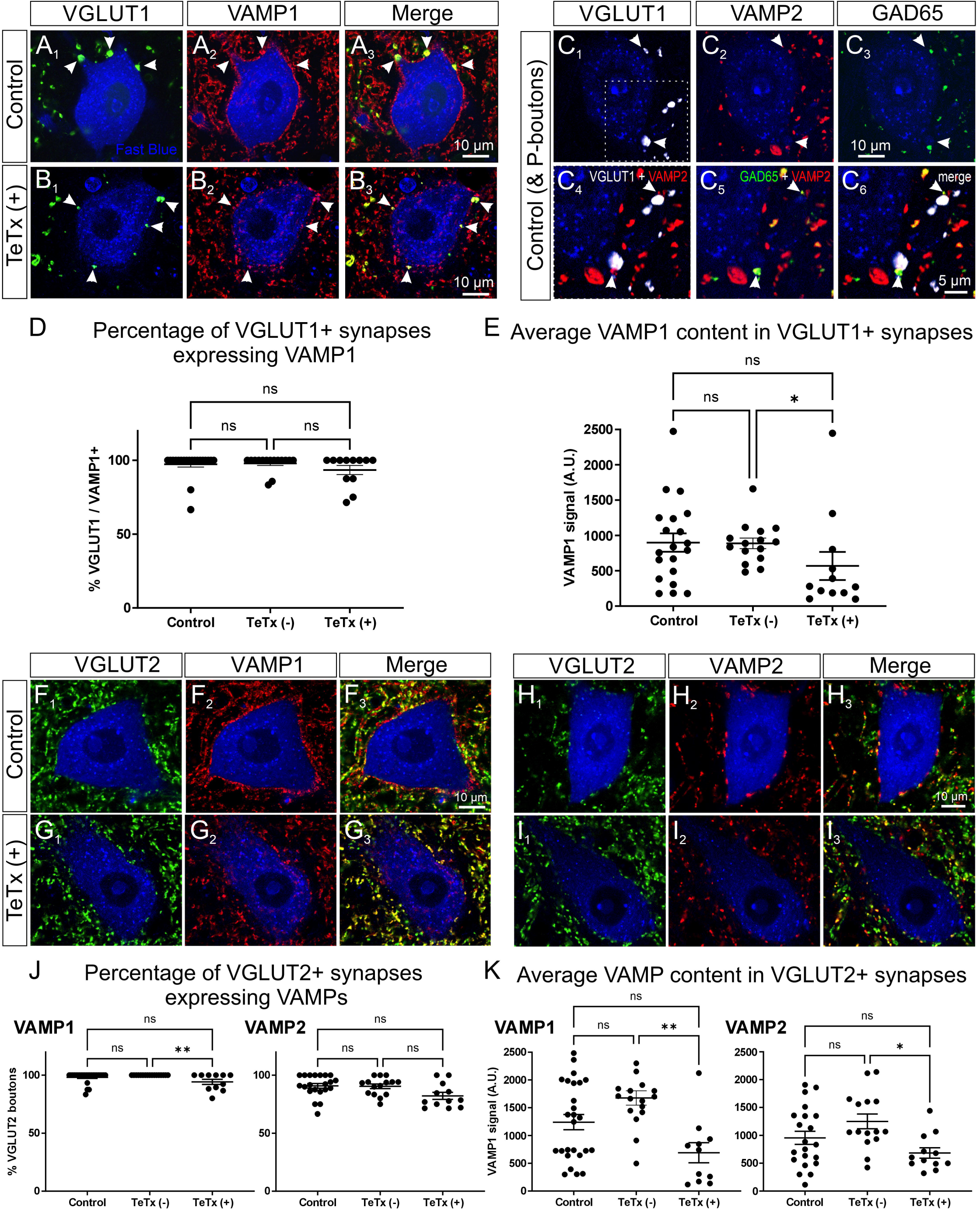
Tetanus toxin effects on VAMP content inside excitatory synapses on TA motoneurons cell bodies. **A1-3,** VAMP1 (red, Cy3) inside VGLUT1 synapses (green, FITC) over control TA motoneurons labeled with Fast Blue. Arrowheads indicate co-localization. **B1-3,** Similar images in animals in which the TA muscle was injected with TeTx and showed positive tetanization 7 days after and just prior collecting tissues for analysis. **C1-6,** Quadruple fluorescent images showing VAMP2 (red, Cy3), VGLUT1 (white, Cy5) and GAD65 (green, FITC) on control TA motoneurons. VAMP2 is not inside VGLUT1 synapses but surrounds and overlaps them because VAMP2 is present in GAD65 P-boutons (arrowheads). **D,** Percentage of VGLUT1 synapses with VAMP1 content (estimated as 2 standard deviations above average background) in controls or animals injected with TeTx in the TA and that showed no (TeTx(-)) or positive tetanization (TeTx(+)). In all graphs each dot represents one motoneuron and average and standard deviations (error bars) are indicated. No significant differences were detected; one-way Kruskal-Wallis followed by Dunn’s multiple comparisons. Statistics details in supplemental table 4. **E,** VAMP1 content in VGLUT1 synapses shows a small depletion in VAMP1 inside VGLUT1 synapses on tetanized TA motoneurons. One-way Kruskal-Wallis, p=0.0393. Post-hoc Dunn’s tests detected significance only between TeTx(-) and TeTx(+) groups (*p=0.0470). Statistics details in Supplementary Table 5. **F1- 3** and **G1-3** VAMP1 in VGLUT2 synapses in Control (**F**) and TeTx(+) TA motoneurons. (**G**). **H1-3** and **I1-3** VAMP2 in VGLUT2 synapses in Control (**F**) and TeTx(+) (**G**) TA motoneurons. **J,** VGLUT2 synapse percentages showing VAMP1 or VAMP2 in TeTx(-), TeTx(+) or controls animals. Small differences were detected between TeTx(-) and TeTx(+) animals. Kruskal-Wallis comparison: p=0.0078 for VAMP1 and p=0.0495 for VAMP2. Post-hoc Dunn’s tests show significance only between TeTx(-) and TeTx(+) for VAMP1. Statistical details in supplementary table 6. **K,** VAMP1 and VAMP2 content inside VGLUT2 synapses on TA motoneurons. Differences in both VAMP1 and VAMP2 were detected comparing TeTx(-) and TeTx(+) animals. Kruskal-Wallis: p=0.0034 for VAMP1 and p=0.0194 for VAMP2. Post-hoc Dunn’s tests demonstrated significance between TeTx(-) and TeTx(+) for VAMP1 and VAMP2. Statistical details in supplementary Table 7. In all graphs *p<0.01 and **p<0.001. *VAMP1 and VAMP2, Vesicle-associated membrane protein 1 and 2; VGLUT1: Vesicular glutamate transporter 1; VGLUT2: Vesicular glutamate transporter 2; GAD65: Glutamic Acid Decarboxylase of 65 kDa: TeTx: Tetanus Toxin; Cy5: Cyanine5; Cy3: Cyanine3; FITC, Fluorescein isothiocyanate; TA: Tibialis Anterior*.

**Figure 6.**
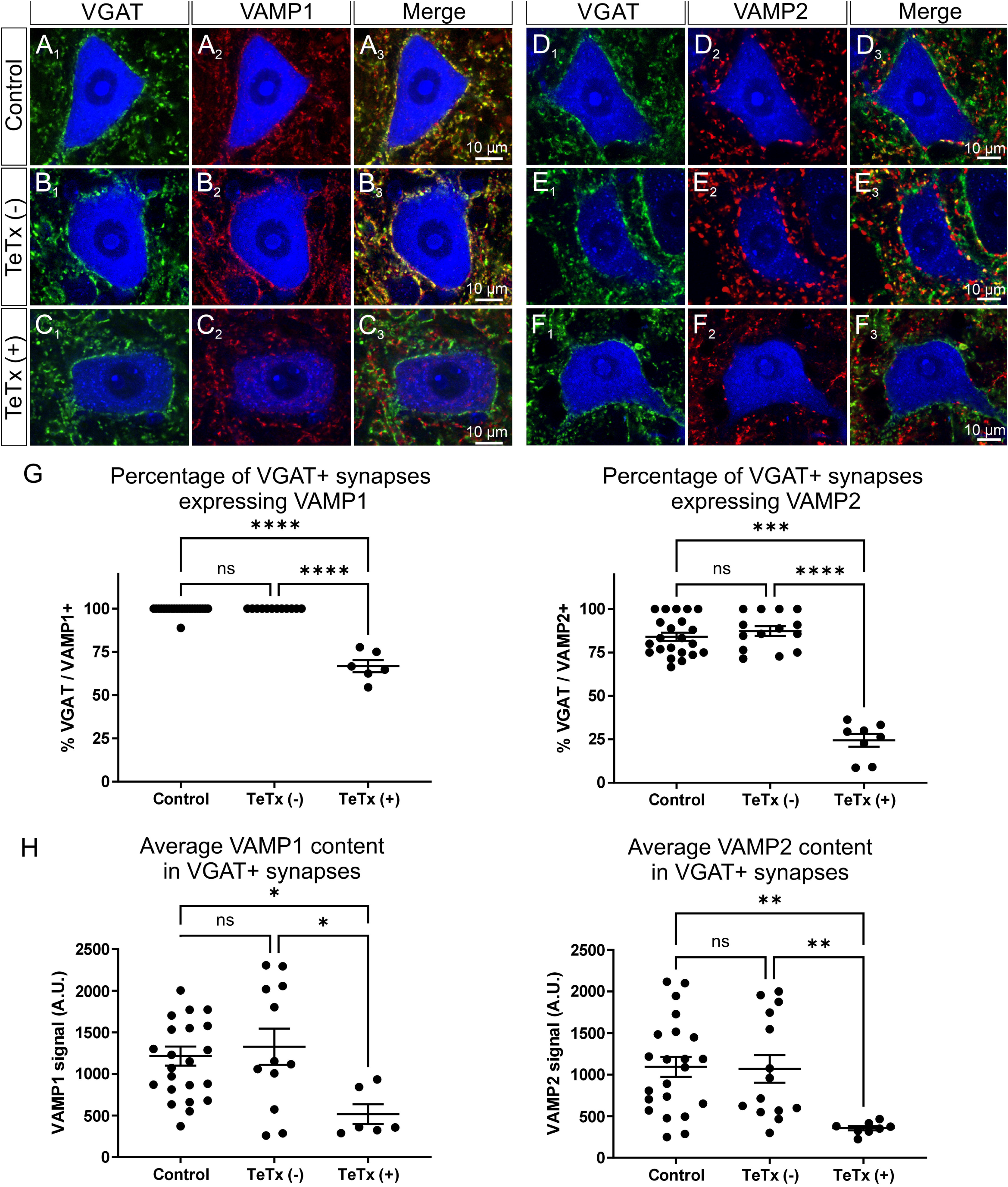
Tetanus toxin effects on VAMP content inside inhibitory synapses on TA motoneurons cell bodies. **A1-3** VAMP1 (red, Cy3) content inside inhibitory synapses (VGAT, green, FITC) on the cell bodies of Fast Blue retrogradely labeled TA motoneurons. **B1-3** VAMP1 content inside VGAT synapses on TA motoneurons treated with TeTx, but not showing tetanization in the TA muscle (TeTx(-)). **B1-3** VAMP1 content inside VGAT synapses on TA motoneurons treated with TeTx and showing positive tetanization in the TA muscle (TeTx(+)). **D1-3**, **E1-3**, and **F1-3**, Similar image composition, as in A1-3, B1-3 and C1-3, but for VAMP2. **G**, Percentage of VGAT synapses containing VAMP1 (graph at left) or VAMP2 (graph at right). A significant depletion for both VAMPs was observed in motoneurons from animals in which the TA injection of TeTx was effective (TeTx(+)). These were significant different to control and TeTx(-) animals. Kruskal-Wallis showed very significant differences for both VAMP1 and VAMP2 (both p<0.0001) and Dunn’s pair-wise multiple tests showed significant differences: ***p<0.001, ****p<0.0001. Statistics details in supplemental table 8. In all graphs each dot represents one motoneuron and average and standard deviations (error bars) are indicated. **H,** VAMP1 (left graph) and VAMP2 (right graph) inside VGAT synapse positive for each VAMP. The TeTx group showed significant differences when compared to the control or the TeTx(-). Kruskal-Wallis showed very significant differences form both VAMP1 (p=0.0185) and VAMP2 ( p=0.0010) and Dunn’s pair-wise multiple tests showed significant differences: *p<0.05, **p<0.01. Stats details in supplemental table 9. *VAMP1 and VAMP2, Vesicle-associated membrane protein 1 and 2; VGAT, Vesicular GABA/Glycine transporter; TeTx: Tetanus Toxin; Cy3: Cyanine3; FITC, Fluorescein isothiocyanate; TA: Tibialis Anterior*.

For quantification, we threshold all VGLUT1, VGLUT2 and VGAT synapses surrounding Fast Blue labeled TA motoneurons and individualize each synaptic bouton. Then we measured VAMP immunofluorescence intensity within the bouton and compared it to average background labeling. Background labeling was obtained by tracing 4 square regions of interest (ROIs) of 3X3 µm inside the Fast Blue labeled nucleolus in the same confocal planes. Any synaptic bouton with VAMP1 or VAMP2 immunofluorescence intensity higher than average ROI labeling +2 standard deviations was considered positive. We pooled together all motoneurons sampled from different animals in each experimental group (data structure shown in Supplementary Table 10) and estimated the percentage of synaptic terminals with VAMP1 or VAMP2 content. In control TA motoneurons, VAMP1 was present in 96.1% ±9.8 (±SD) of VGLUT1 synapses (n=20 motoneurons; 114 synapses), 98.1% ±4.7 of VGLUT2 synapses (n=26 motoneurons; 374 synapses) and 99.5% ±2.3 of VGAT synapses (n=23 motoneurons, 329 synapses), while VAMP2 was detected in 90.8% ±9.4 and 84.1% ±11.1 of respectively VGLUT2 and VGAT synapses (n=21 and 23 motoneurons, 316 and 299 synapses). The percentages of VAMP1 and VAMP2 labeled synapses were not significantly different in TA motoneurons innervating muscles that showed little or no tetanization (Figures 5D, 5J and 5G; Supplemental Tables 4 and 6). TA motoneurons innervating tetanized muscles showed no change in the percentage of VGLUT1 synapses positively labeled for VAMP1 (93.5% ±10.6; n=12 motoneurons) (Figure 5D). The percentage of VGLUT2 synapses positive for VAMP1 or VAMP2 showed small decreases (VAMP1: 94.2% ±7.2, n=12 motoneurons, 66 synapses; 82.2% ±10.5, n=12 motoneurons, 187 synapses) but these did not reach significance compared to control (VAMP1, p=0.081; VAMP2 p=0.057; Dunn’s test to control following Kruskal-Wallis ANOVA) (Figure 5J). In contrast, large and significant changes occurred on inhibitory synapses (Figure 6G, Supplemental Table 8). The percentage of VGAT synapses expressing VAMP1 dropped to 66.9% ±8.5 (n=6 motoneurons, 87 synapses; p<0.0001 Dunn’s test to control) and the percentage expressing VAMP2 to 24.5% ±10.5 (n=8 motoneurons, 149 synapses; p<0.0001 Dunn’s test to control following a Kruskal-Wallis ANOVA). For analyses of labeling intensity, we only considered positively labeled synapses (2 standard deviations above average background). VAMP1 immunofluorescence was reduced by approximately 36% and 44% from control values in respectively VGLUT1 and VGLUT2 labeled synapses, and VAMP2 was reduced 28.4% in VGLUT2 synapses. These depletions although large did not reach significance compared to controls (Dunn’s tests p=0.1161, 0.1443 and 0.4239 for Hedges g estimates of 0.5, 0.8 and 0.6; Supplemental Tables 5 and 7), but did reach significance when compared to TA motoneurons with no tetanization after TeTx injections (Figures 5E and 5K). In comparison to the mild effects observed in VGLUT1 and VGLUT2 synapses, the decrease in VAMP signals was much larger and significant in VGAT synapses compared to either controls or TeTx treated showing little to no tetanization (Figures 6G and 6H; Supplemental Table 9). Compared to controls, VAMP1 signals were reduced by 59% (p=0.0274, Dunn’s test, g=1.5) and VAMP2 by 67% (p=0.001, Dunn’s test. G=1.5) in synapses that remained positive for VAMP.

In conclusion, TeTx injected in muscle reduces VAMP1 and VAMP2 content very significantly in inhibitory synapses surrounding the motoneuron cell body, but the effects of the toxin are much milder on excitatory synapses. Moreover, animals injected with TeTx which did not show strong tetanization displayed none or only small reductions in VAMP content. We conclude that the functional effect of TA tetanization is due to removal of VAMPs and consequent decreased of neurotransmitter release largely from inhibitory synapses.

## Discussion

The results demonstrate that transient disruption of GABA and/or glycine neurotransmitter release on axotomized motoneurons for around one week after injury using mild doses of tetanus toxin (TeTx) significantly delays muscle reinnervation. We interpret this result to suggest that these neurotransmitters contribute to the regeneration phenotype of motoneurons while growing back their axons towards muscles after nerve injuries. Moreover, these synapses might be key facilitators of activity-dependent mechanisms promoting nerve regeneration (Gordon *et al*., 2009; Gordon, 2016; Gordon & English, 2016). This finding opens a new window for possible therapeutics to speed motor axon regeneration and diminish detrimental effects of chronic denervation and for possibly extending regenerative capacity in motoneurons when long periods of regeneration or delayed nerve repairs are required.

Synaptically released GABA and/or glycine acting on GABA_A_ and glycine receptors on the surface of axotomized spinal motoneurons are predicted to induce outward chloride currents that depolarize the neuron in the absence of KCC2 (Akhter *et al*., 2019; Akhter, 2020). This was previously shown on brainstem axotomized motoneurons of the dorsal nucleus of the vagus (Nabekura *et al*., 2002) and the facial nucleus (Toyoda *et al*., 2003). Injured facial motoneurons showed calcium transients due to the opening of voltage-gated calcium channels driven by depolarizations dependent on GABA_A_ and NMDA receptor activation. Potentially these calcium signals at the level of the cell body can facilitate or maintain expression of the regenerative phenotype acting on the many regeneration associated genes (RAGs) with calcium dependent promoters. Hundreds of RAGs are expressed after axotomy forming a complex gene network organized by a few hub proteins upregulated immediately after axotomy and that include the transcription factors ATF3 and Jun (Stam *et al*., 2007; Michaelevski *et al*., 2010; Chandran *et al*., 2016). RAGs that depend on Jun activation use activating-protein-1 (AP1) like sequences (Mason *et al*., 2022), while those responsive to ATF3 or ATF3-Jun dimers use cAMP-responsive element CRE-like sequences that require cAMP and also importantly calcium (Gey *et al*., 2016). The regulation of these late RAG networks coincides temporally with the time course of KCC2 downregulation; the *kcc2* gene completely shuts off by 3 days after injury while KCC2 protein is progressively removed from the membrane during the first week post-injury and peaking depletion in the second week (Akhter *et al*., 2019). Our interventions blocked synaptic release of GABA and glycine with TeTx during the first week post-axotomy, a period characterized by axon sprouting, bridging the injury site and entering the distal stump initiating axon growth (Gordon & Borschel, 2017; Wariyar *et al*., 2022). Slowing these initial regenerative mechanisms likely delayed muscle reinnervation, as shown in our study.

Removal of KCC2 is critical in this potential mechanism and this is supported by two recent studies, although both have interpretative caveats that need careful consideration. In one study, KCC2 downregulation in axotomized motoneurons was prevented by upregulating expression of a *kcc2* transgene depending on CamKII upregulation in injured motoneurons (Cheung *et al*., 2023). KCC2 maintenance correlated with impaired post-injury recovery of motor function assessed via an accelerating rotarod and sciatic static index estimates. Although this conclusion supports ours, there are alternative interpretations to the study. The motor behaviors measured correlate with nerve regeneration only poorly and could be affected by KCC2 overexpression in many other CamKII-expressing neurons throughout the brain, including those in several important motor centers like cerebellum and motor cortex. A more recent paper demonstrated enhanced axon regeneration in the peripheral nerves of *kcc2* heterozygotes compared to wild types after tibial nerve injuries (Ando *et al*., 2024). This result also suggests that reduced KCC2 accelerates axon regeneration, however interpretation is constrained by the fact that *kcc2* gene expression is expected to be completely shut down in injured motoneurons in both wild-types and *kcc2* heterozygotes. Thus, the results might be best explained by faster downregulation of KCC2 in heterozygotes with maybe lower initial content of KCC2 protein. Unfortunately, temporal course of KCC2 levels in axotomized motoneurons of wild-types and heterozygotes was not directly compared. Our study shows more directly the need of GABA and/or glycine release to promote axon regeneration in axotomized motoneurons. It supports conclusions advanced in these recent papers, despite the possibility of alternative explanations in each of them.

A critical role of GABA/glycinergic synapses in axon regeneration is also supported by studies of synaptic plasticity on axotomized motoneurons. It is well known that after axotomy the cell body of motoneurons is “stripped” of synapses but with differential mechanisms on inhibitory vs excitatory synapses resulting in comparatively more preservation of inhibitory synapses (Alvarez *et al*., 2020). Remarkably, enhancing “inhibitory” synapses maintenance after axotomy always correlated with faster recovery of motor function. This was the case when reduced removal of inhibitory synapses occurred in transgenic animals with absent complement C3 (Berg *et al*., 2012), Major Histocompatibility Complex I (MHCI) (Oliveira *et al*., 2004) or cadherin epidermal growth factor (EGF) laminin G (LAG) seven-pass G-type receptor 2 (Celrs 2) (Yu *et al*., 2023). Similarly, this was also the case following selective preservation of inhibitory synapses via exogenous application of BDNF (Novikov *et al*., 2000). The fact that these disparate animal models all point to a correlation between inhibitory synapse preservation and motor axon regeneration supports our conclusion that GABA and/or glycine exert faciliatory roles on regeneration. Many of these papers interpreted their results within the standard contemporary consensus in the field for a necessary alteration of E/I ratios towards enhanced inhibition to redirect metabolic support towards protein synthesis and regeneration. However, it is now well established that “inhibitory” synapses on axotomized motoneurons more likely provide some excitatory drive.

Our results also align with studies that investigated the role of trophic factors on activity-dependent facilitation of axon regeneration and revealed the significance of BDNF/TrkB signaling (Al-Majed *et al*., 2000a; McGregor & English, 2018). Initial studies showed that BDNF promotes the growth of cut axons into nerve grafts (Zhang et al., 2000; English et al., 2005) and that BDNF is critically upregulated after nerve injury in motoneurons, Schwann cells and many subtypes of sensory neurons, but with each cell type following different time courses (Meyer *et al*., 1992; Funakoshi *et al*., 1993; Gordon, 2009). Motoneurons upregulate BDNF shortly after axotomy, but this is transient and only lasts for a few days (Kobayashi *et al*., 1996; Al-Majed *et al*., 2000a), while upregulation in peripheral Schwann cells is slower but maintained for weeks (Funakoshi *et al*., 1993; Zhang *et al*., 2000). Nonetheless, cell-specific deletion studies demonstrated that BDNF expression and TrkB signaling in motoneurons are critically important for exercise promotion of regeneration (English *et al*., 2007; Wilhelm *et al*., 2012). BDNF could help maintain inhibitory synapses (Novikov *et al*., 2000) and although we found that endogenous BDNF/TrkB signaling is not necessary to downregulate KCC2 in motoneurons after axotomy (Akhter *et al*., 2017), more recently we reported that exogenous BDNF accelerates KCC2 removal (Capilla-Lopez *et al*., 2024). This can be explained by considering KCC2 removal as a two-step process in which gene expression is suppressed first and then membrane KCC2 is removed. BDNF appears to act on trafficking and turn-over of membrane KCC2 in axotomized motoneurons but has no action of *kcc2* gene expression. Thus, while BDNF may accelerate removal, blocking BDNF cannot prevent the shutdown of the gene and the eventual loss of membrane KCC2 by normal turn-over without replenishment. The time course of BDNF expression in motoneurons is well fitted to accelerate removal of KCC2 soon after injury and facilitate the transition of GABA and glycine actions to depolarizing. Long-term BDNF actions on KCC2 membrane trafficking are irrelevant without *de novo* upregulation of the *kcc2* gene. Several authors, including our group, have reported that *kcc2* gene expression recovers when motor axons reinnervate muscle (Tatetsu *et al*., 2012; Kim *et al*., 2018; Akhter *et al*., 2019). This suggests that signals from muscle upregulate *kcc2* gene expression when regeneration is completed, however it does not rule out alternative mechanisms for *kcc2* upregulation and loss of GABA/glycine facilitation of regenerative mechanisms during a prolonged regeneration period and delayed muscle reinnervation. Our experimental animals have relatively small sizes and therefore the period of axon regeneration before reaching muscle targets is short compared to larger species, including humans. Whether recovery of KCC2 expression, affecting GABA and/or glycine actions on regeneration, occurs during extended periods of regeneration is yet untested, though this knowledge remains important since it might help explaining mechanisms that decrease motoneuron regenerative capacity with time after injury (Fu & Gordon, 1995b; a).

### Limitations of the study and future studies.

Our experimental design blocked GABA and/or glycine release only for a brief period after injury. This was unavoidable because we used small doses of TeTx to prevent systemic effects or pleiotropic actions in other muscles that might obscure interpretation of any results. Whether more complete and long-lasting blocks of GABAergic and glycinergic synapses will cause more profound delays or abolition of functional regeneration cannot be determined at present. This affects interpretation of the results on whether GABA and glycine are facilitators that augment and speed-up regeneration or whether they are necessary for the regenerative phenotype. Moreover, the long-term action of GABA and glycine during delayed regeneration remains an open question. Future studies should address these key issues by genetic manipulation of KCC2 levels in specifically axotomized motoneurons with better control of timing and cell specificity than has been yet possible.

A second limitation is regarding the translatability of the phenomenon we describe. It is unlikely that global pharmacological approaches to enhance GABA neurotransmission would be helpful in the clinic giving the many side effects of these drugs. One aspect that was not studied here is the exact inhibitory circuits that are the origin of the GABA/glycinergic synapses retained by axotomized motoneurons and responsible for the effects we described. Exact knowledge of these interneurons is critical for future studies to activate these circuits specifically over motoneurons that are regenerating. This could be accomplished taking advantage of reflex pathways and their accessibility to activation from the periphery. Moreover, knowledge of the exact interneurons which facilitate a regenerative phenotype in motoneurons could lead the way to new experimental approaches in animal models for future genetic dissection with the purpose of manipulating activity or synaptic release and address questions on their necessity and actions in acute or delayed regeneration paradigms.

## Acknowledgments

This work was supported by NIH grants R01 NS111969 and R21 NS114839 to FJA. PMC was the recipient of a postdoctoral Margarita Salas award from the University of Seville/EU. We also acknowledge William Goolsby, Director of the Cell Biology Electronics and Mechanical Department Core for his contribution building the pressure probe used in these studies and Olivia Mistretta for her help in the initial titer of toxin batches.

## Author Contributions

Conceptualization, FJA.

Methodology, RLW, PMC, WMM, AWE, and FJA.

Formal analysis, RLW, PMC, and FJA,

Investigation, RLW, PMC, WMM, AWE, and FJA.

Resources, FJA and AEW.

Data curation and Figures, RLW, PMC and FJA.

Writing –original draft, FJA.

Writing – review & editing, RLW, PMC, AWE, and FJA.

Supervision, FJA and AEW.

Project administration, FJA.

Funding acquisition, FJA.

## Data Accessibility

The data files of this manuscript will be uploaded to the online submission system to be deposited with Figshare.

All statistical analyses details are reported in the Supplemental Information file as supplementary statistical tables 1 to 10.

Any additional confocal images can be shared with readers of the manuscript upon request to the corresponding author.

## ABBREVIATIONS

TeTx: Tetanus Toxin
TA: Tibialis Anterior Muscle
LG: Lateral Gastrocnemius Muscle
GABA: Gamma-aminobutyric acid
KCC2: Potassium chloride cotransporter 2
VAMP: Vesicle-associated membrane protein
AVMA: American Veterinary Medical Association
PP: Pressure-Probe
M-response: Muscle response
EMG: Electromyography
CMAP: Compound Muscle Action Potential
VGLUT1: Vesicular glutamate transporter 1
VGLUT2: Vesicular glutamate transporter 2
VGAT: Vesicular GABA/Glycine transporter (also known as VIAAT, Vesicular inhibitory amino acid transporter)
VACHT: Vesicular acetylcholine transporter
NeuN: Neuronal Nucleus marker
GAD65: Glutamic Acid Decarboxylase of 65 kDa
NMJ: Neuromuscular Junction
NFH: Neurofilament Heavy subunit
ANOVA: Analysis of Variance
Cy5: Cyanine5
Cy3: Cyanine3
FITC: Fluorescein isothiocyanate
NMDA: N-methyl-D-aspartate
RAG: regeneration associated gene
ATF3: Activating transcription factor 3
AP1: Activating protein 1
MHCI: Major Histocompatibility
Celrs: Complex I cadherin EGF LAG seven-pass G-type receptor 2
BDNF: Brain Derived Neurotrophic Factor
TrkB: tyrosine protein kinase B
P-bouton: synaptic boutons presynaptic to primary afferent synapses

