## Supplemental Figures and Tables for "GABA and glycine synaptic release on axotomized motoneuron cell bodies promotes motor axon regeneration"

**Supplementary Table 1 (for Figure 3D).**

**Average M-response in TA muscles**

Two-Way ANOVA for time after injury and experimental group:

Sham, Tetanus, Crush and Crush + Tetanus

- Time after Injury: F_(3, 94)_ = 19.42	p < 0.0001
- Experimental group: F_(3, 94)_ = 45.57	p < 0.0001
- Interaction: F_(9, 94)_ = 7.197	p < 0.0001

| Tukey's multiple comparisons test | Predicted mean diff. | 95% CI of diff. | % 2^nd^ vs 1^st^ difference | Hedge’s g | q | DF | Adjusted P Value | N1 | N2 |
| --- | --- | --- | --- | --- | --- | --- | --- | --- | --- |
| **0 days post-injury** |  |  |  |  |  |  |  |  |  |
| Sham vs. TeTx | -0.1346 | -0.4228 to 0.1537 | 11.9 | -0.5 | 1.805 | 27.39 | 0.5852 | 17 | 13 |
| Sham vs. Crush | 0.02947 | -0.2680 to 0.3269 | -2.7 | 0.1 | 0.3836 | 26.84 | 0.9929 | 17 | 13 |
| Sham vs. TeTx + Crush | 0.05367 | -0.2466 to 0.3540 | -4.8 | 0.2 | 0.7053 | 20.68 | 0.9584 | 17 | 9 |
| TeTx vs. Crush | 0.1640 | -0.1364 to 0.4645 | -13.0 | 0.6 | 2.131 | 23.92 | 0.4494 | 13 | 13 |
| TeTx vs. TeTx + Crush | 0.1882 | -0.1152 to 0.4917 | -14.9 | 0.7 | 2.468 | 18.87 | 0.3292 | 13 | 9 |
| Crush vs. TeTx + Crush | 0.02420 | -0.2874 to 0.3358 | -2.2 | 0.1 | 0.3084 | 19.34 | 0.9962 | 13 | 9 |
| **10 days post-injury** |  |  |  |  |  |  |  |  |  |
| Sham vs. TeTx | -0.03879 | -0.6072 to 0.5296 | 3.4 | -0.1 | 0.3130 | 7.584 | 0.9958 | 5 | 5 |
| Sham vs. Crush | 1.064 | 0.5104 to 1.618 | **-92.3** | **4.9** | **10.78** | **4.216** | **0.0045**** | 5 | 3 |
| Sham vs. TeTx + Crush | 1.048 | 0.4978 to 1.599 | **-90.9** | **4.1** | **10.54** | **4.344** | **0.0045**** | 5 | 3 |
| TeTx vs. Crush | 1.103 | 0.6692 to 1.537 | **-92.5** | **5.5** | **14.08** | **4.344** | **0.0014**** | 5 | 3 |
| TeTx vs. TeTx + Crush | 1.087 | 0.6565 to 1.518 | **-91.2** | **5.4** | **13.70** | **4.544** | **0.0012**** | 5 | 3 |
| Crush vs. TeTx + Crush | -0.01575 | -0.1699 to 0.1384 | 17.6 | -0.3 | 0.6044 | 3.792 | 0.9705 | 3 | 3 |
| **20 days post-injury** |  |  |  |  |  |  |  |  |  |
| Sham vs. TeTx | 0.09855 | -0.5091 to 0.7062 | -8.0 | 0.4 | 0.8801 | 4.543 | 0.9203 | 6 | 4 |
| Sham vs. Crush | 0.6469 | 0.3325 to 0.9614 | **-52.4** | **4.0** | **9.391** | **7.735** | **0.0008***** | 6 | 4 |
| Sham vs. TeTx + Crush | 0.9153 | 0.6290 to 1.202 | **-74.1** | **5.7** | **13.91** | **9.708** | **<0.0001****** | 6 | 6 |
| TeTx vs. Crush | 0.5484 | -0.06845 to 1.165 | *-48.2* | **2.5** | *4.964* | *4.262* | *0.0732* | 4 | 4 |
| TeTx vs. TeTx + Crush | 0.8168 | 0.1987 to 1.435 | **-71.8** | **3.9** | **7.523** | **4.092** | **0.0193*** | 4 | 6 |
| Crush vs. TeTx + Crush | 0.2684 | -0.0268 to 0.5636 | *-45.6* | **1.9** | *4.247* | *7.058* | *0.0741* | 4 | 6 |
| **30 days post-injury** |  |  |  |  |  |  |  |  |  |
| Sham vs. TeTx | -0.06511 | -0.3552 to 0.2250 | 5.6 | -0.5 | 1.065 | 6.671 | 0.8727 | 6 | 4 |
| Sham vs. Crush | 0.2132 | -0.2683 to 0.6947 | -18.5 | 0.8 | 2.099 | 6.689 | 0.4952 | 6 | 6 |
| Sham vs. TeTx + Crush | 0.7022 | 0.4940 to 0.9104 | **-60.9** | **6.1** | **15.03** | **8.612** | **<0.0001****** | 6 | 6 |
| TeTx vs. Crush | 0.2783 | -0.2111 to 0.7678 | -22.9 | 1.0 | 2.653 | 7.085 | 0.3168 | 4 | 6 |
| TeTx vs. TeTx + Crush | 0.7673 | 0.4831 to 1.052 | **-63.0** | **7.1** | **14.34** | **4.784** | **0.0008***** | 4 | 6 |
| Crush vs. TeTx + Crush | 0.4890 | 0.0063 to 0.9717 | **-52.0** | **2.1** | **5.032** | **5.739** | **0.0475*** | 6 | 6 |

**Supplementary Table 2 (for Figure 3E).**

**Average M-response in LG muscles**

Two-Way ANOVA for time after injury and experimental group:

Sham, Tetanus, Crush and Crush + Tetanus

- Time after Injury: F_(3, 40)_ = 24.57	p < 0.0001
- Experimental group: F_(3, 54)_ = 24.00	p < 0.0001
- Interaction: F_(9, 40)_ = 10.21	p < 0.0001

| Tukey's multiple comparisons test | Predicted mean diff. | 95% CI of diff. | % 2^nd^ vs 1^st^ difference | Hedge’s  g | q | DF | Adjusted P Value | N1 | N2 |
| --- | --- | --- | --- | --- | --- | --- | --- | --- | --- |
| **0 days post-injury** |  |  |  |  |  |  |  |  |  |
| Sham vs. TeTx | 0.06529 | -0.1418 to 0.2723 | -6.0 | 0.3 | 1.221 | 26.67 | 0.8234 | 17 | 13 |
| Sham vs. Crush | -0.03104 | -0.3251 to 0.2630 | 2.9 | -0.1 | 0.4191 | 19.34 | 0.9907 | 17 | 13 |
| Sham vs. TeTx + Crush | 0.001623 | -0.2706 to 0.2738 | -0.2 | 0.0 | 0.02434 | 14.84 | >0.9999 | 17 | 9 |
| TeTx vs. Crush | -0.09633 | -0.3957 to 0.2030 | 9.5 | -0.4 | 1.275 | 19.80 | 0.8041 | 13 | 13 |
| TeTx vs. TeTx + Crush | -0.06366 | -0.3415 to 0.2142 | 6.2 | -0.3 | 0.9316 | 15.33 | 0.9109 | 13 | 9 |
| Crush vs. TeTx + Crush | 0.03267 | -0.3056 to 0.3709 | -3.0 | 0.1 | 0.3824 | 19.93 | 0.9929 | 13 | 9 |
| **10 days post-injury** |  |  |  |  |  |  |  |  |  |
| Sham vs. TeTx | 0.03835 | -0.5556 to 0.6323 | -3.6 | 0.1 | 0.2929 | 7.940 | 0.9966 | 5 | 5 |
| Sham vs. Crush | 0.9524 | 0.4596 to 1.445 | **-88.1** | **4.7** | **10.04** | **5.061** | **0.0031**** | 5 | 3 |
| Sham vs. TeTx + Crush | 0.9298 | 0.4355 to 1.424 | **-86.0** | **4.0** | **10.01** | **4.764** | **0.0040**** | 5 | 3 |
| TeTx vs. Crush | 0.9141 | 0.3761 to 1.452 | **-87.7** | **3.6** | **8.926** | **4.915** | **0.0059**** | 5 | 3 |
| TeTx vs. TeTx + Crush | 0.8914 | 0.3510 to 1.432 | **-85.5** | **3.5** | **8.862** | **4.652** | **0.0072**** | 5 | 3 |
| Crush vs. TeTx + Crush | -0.02262 | -0.2832 to 0.2379 | 17.6 | -0.3 | 0.5086 | 3.866 | 0.9819 | 3 | 3 |
| **20 days post-injury** |  |  |  |  |  |  |  |  |  |
| Sham vs. TeTx | 0.1407 | -0.4255 to 0.7069 | -12.3 | 0.6 | 1.194 | 6.380 | 0.8322 | 6 | 4 |
| Sham vs. Crush | 0.6728 | 0.2613 to 1.084 | **-58.6** | **3.1** | **7.406** | **7.991** | **0.0035**** | 6 | 4 |
| Sham vs. TeTx + Crush | 0.7204 | 0.3437 to 1.097 | **-62.7** | **3.6** | **8.796** | **7.495** | **0.0015**** | 6 | 6 |
| TeTx vs. Crush | 0.5321 | -0.03574 to 1.100 | *-52.8* | **2.5** | *4.950* | *4.848* | *0.0629* | 4 | 4 |
| TeTx vs. TeTx + Crush | 0.5797 | 0.00393 to 1.155 | **-57.5** | **3.0** | **5.794** | **4.004** | **0.0489**** | 4 | 6 |
| Crush vs. TeTx + Crush | 0.04755 | -0.2806 to 0.3757 | -10.0 | 0.3 | 0.7188 | 5.762 | 0.9540 | 4 | 6 |
| **30 days post-injury** |  |  |  |  |  |  |  |  |  |
| Sham vs. TeTx | -0.02899 | -0.3546 to 0.2967 | 2.6 | -0.2 | 0.4392 | 5.858 | 0.9886 | 6 | 4 |
| Sham vs. Crush | 0.2140 | -0.1857 to 0.6138 | -19.1 | 0.9 | 2.481 | 7.282 | 0.3643 | 6 | 6 |
| Sham vs. TeTx + Crush | 0.3337 | 0.0097 to 0.6577 | **-29.8** | **1.9** | **4.620** | **8.348** | **0.0436*** | 6 | 6 |
| TeTx vs. Crush | 0.2430 | -0.1853 to 0.6714 | -21.2 | 1.0 | 2.576 | 7.921 | 0.3316 | 4 | 6 |
| TeTx vs. TeTx + Crush | 0.3627 | -0.0087 to 0.7342 | *-31.6* | **1.9** | *4.438* | *7.876* | *0.0556* | 4 | 6 |
| Crush vs. TeTx + Crush | 0.1197 | -0.3119 to 0.5513 | -13.2 | 0.5 | 1.211 | 9.506 | 0.8266 | 6 | 6 |

**Supplementary Table 3 (for Figure 4C).**

**Percentage of reinnervated NMJs in TA and LG**

One-Way ANOVA for experimental group:

Sham, Tetanus, Crush and Crush + Tetanus

- **TA**: F_(3, 19)_ = 9.99997	p = 0.0004
- **LG**: F_(3, 20)_ = 1.617	p = 0.2171

| Tukey's multiple comparisons test | Predicted mean diff. | 95% CI of diff. | % 2^nd^ vs 1^st^ difference | Hedge’s  g | q | DF | Adjusted P Value | N1 | N2 |
| --- | --- | --- | --- | --- | --- | --- | --- | --- | --- |
| **Tibialis Anterior** |  |  |  |  |  |  |  |  |  |
| Sham vs. TeTx | 0.6857 | -4.733 to 6.104 | -0.8 | 0.2 | 0.5032 | 19 | 0.9841 | 6 | 5 |
| Sham vs. Crush | 2.368 | -2.798 to 7.535 | -2.7 | 0.8 | 1.823 | 19 | 0.5806 | 6 | 6 |
| Sham vs. TeTx + Crush | 9.068 | 3.902 to 14.23 | **-10.3** | **2.8** | **6.980** | **19** | **0.0005***** | 6 | 6 |
| TeTx vs. Crush | 1.683 | -3.736 to 7.101 | -1.9 | 0.5 | 1.235 | 19 | 0.8185 | 5 | 6 |
| TeTx vs. TeTx + Crush | 8.383 | 2.964 to 13.80 | **-9.6** | **2.3** | **6.152** | **19** | **0.0018**** | 5 | 6 |
| Crush vs. TeTx + Crush | 6.700 | 1.534 to 11.87 | **-7.8** | **2.2** | **5.157** | **19** | **0.0085**** | 6 | 6 |
| **Lateral Gastrocnemius** |  |  |  |  |  |  |  |  |  |
| Sham vs. TeTx | -0.7030 | -5.622 to 4.216 | 0.8 | -0.2 | 0.5657 | 20 | 0.9777 | 6 | 5 |
| Sham vs. Crush | 1.258 | -3.432 to 5.949 | -1.4 | 0.4 | 1.062 | 20 | 0.8752 | 6 | 6 |
| Sham vs. TeTx + Crush | 2.691 | -1.829 to 7.210 | -3.1 | 1.0 | 2.357 | 20 | 0.3666 | 6 | 7 |
| TeTx vs. Crush | 1.961 | -2.958 to 6.881 | -2.2 | 0.6 | 1.578 | 20 | 0.6842 | 5 | 6 |
| TeTx vs. TeTx + Crush | 3.394 | -1.363 to 8.151 | -3.8 | 1.3 | 2.824 | 20 | 0.2223 | 5 | 7 |
| Crush vs. TeTx + Crush | 1.432 | -3.087 to 5.952 | -1.6 | 0.6 | 1.254 | 20 | 0.8116 | 6 | 7 |

**Supplementary Table 4 (for Figure 5D).**

**% of VGLUT1+ synapses on TA Fast Blue motoneurons expressing VAMP1**

Kruskal-Wallis test (non-parametric):

p = 0.2580; K-S Statistic = 2.710

Experimental groups:

- Control: non-injected TA,
- TeTx (-): TA injected with TeTx but does not show tetanization,
- TeTx (+): TA injected with TeTx showing positive tetanization.

| Dunn’s multiple comparisons test | mean rank 1 | mean rank 2 | Rank  difference | Mean 1 ± S.D. | Mean 2 ± S.D. | % difference (2-1) | Hedge’s  g | P Value | N1 | N2 | Z |
| --- | --- | --- | --- | --- | --- | --- | --- | --- | --- | --- | --- |
| Control vs TeTx (-) | 25.45 | 25.00 | 0.450 | 96.03% ± 8.49 | 97.94% ± 5.46 | 1.99% | 0.2254 | >0.9999 | 20 | 15 | 0.2695 |
| Control vs TeTx (+) | 25.45 | 20.33 | 5.117 | 96.03% ± 8.49 | 93.45% ± 10.61 | -3.99% | 0.2601 | 0.7976 | 20 | 12 | 1.113 |
| TeTx (-) vs TeTx (+) | 25.00 | 20.33 | 4.667 | 97.94% ± 5.46 | 93.45% ± 10.61 | -4.584 | 0.5517 | 0.5946 | 15 | 12 | 1.287 |

**Supplementary Table 5 (for Figure 5E).**

**Average VAMP1 content in VGLUT1+ synapses on TA Fast Blue motoneurons**

Kruskal-Wallis test (non-parametric):

**p = 0.0393**; K-S Statistic = 6.474

Experimental groups:

- Control: non-injected TA,
- TeTx (-): TA injected with TeTx but does not show tetanization,
- TeTx (+): TA injected with TeTx showing positive tetanization.

| Dunn’s multiple comparisons test | mean rank 1 | mean rank 2 | Rank  difference | Mean 1 ± S.D. | Mean 2 ± S.D. | % difference (2-1) | Hedge’s  g | P Value | N1 | N2 | Z |
| --- | --- | --- | --- | --- | --- | --- | --- | --- | --- | --- | --- |
| Control vs TeTx (-) | 25.85 | 28.33 | -2.483 | 889.8 ± 585.4 | 888.8 ± 291.2 | -0.11% | 0.00207 | >0.9999 | 20 | 15 | 0.5303 |
| Control vs TeTx (+) | 25.85 | 15.50 | 10.35 | 889.8 ± 585.4 | 569.1 ± 687.0 | -36.04% | 0.5135 | 0.1161 | 20 | 12 | 2.067 |
| TeTx (-) vs TeTx (+) | 28.33 | 15.50 | 12.83 | 888.8 ± 291.2 | 569.1 ± 687.0 | **-35.97%** | **0.6329** | **0.0470*** | 15 | 12 | 2.417 |

**Supplementary Table 6 (for Figure 5J).**

**% of VGLUT2+ synapses on TA Fast Blue motoneurons expressing VAMP1 and/or VAMP2**

Kruskal-Wallis test (non-parametric):

VAMP1: **p = 0.0078**; K-S Statistic = 9.704

VAMP2: **p = 0.0495**; K-S Statistic = 6.010

Experimental groups:

- Control: non-injected TA,
- TeTx (-): TA injected with TeTx but does not show tetanization,
- TeTx (+): TA injected with TeTx showing positive tetanization.

| Dunn’s multiple comparisons test | mean rank 1 | mean rank 2 | Rank  difference | Mean 1 ± S.D. | Mean 2 ± S.D. | % difference (2-1) | Hedge’s  g | P Value | N1 | N2 | Z |
| --- | --- | --- | --- | --- | --- | --- | --- | --- | --- | --- | --- |
| **VAMP1** |  |  |  |  |  |  |  |  |  |  |  |
| Control vs TeTx (-) | 27.85 | 32.00 | -4.154 | 98.14% ± 4.67 | 100% ± 0.00 | 1.90% | 0.5100 | 0.5766 | 26 | 17 | 1.304 |
| Control vs TeTx (+) | 27.85 | 19.73 | 8.119 | 98.14% ± 4.67 | 94.24 ± 7.22 | -3.97% | 0.7065 | 0.0812 | 26 | 11 | 2.210 |
| TeTx (-) vs TeTx (+) | 32.00 | 19.73 | 12.27 | 100% ± 0.00 | 94.24 ± 7.22 | **-5.76%** | **1.2864** | **0.0057**** | 17 | 11 | 3.106 |
| **VAMP2** |  |  |  |  |  |  |  |  |  |  |  |
| Control vs TeTx (-) | 27.83 | 26.60 | 1.233 | 90.81% ± 9.44 | 90.44% ± 7.64 | -0.42% | 0.0421 | >0.9999 | 21 | 15 | 0.2628 |
| Control vs TeTx (+) | 27.83 | 16.04 | 11.79 | 90.81% ± 9.44 | 82.19% ± 10.51 | -9.5% | 0.8766 | 0.0567 | 21 | 12 | 2.348 |
| TeTx (-) vs TeTx (+) | 26.60 | 16.04 | 10.56 | 90.44% ± 7.64 | 82.19% ± 10.51 | -9.1% | 0.9150 | 0.1486 | 15 | 12 | 1.964 |

**Supplementary Table 7 (for Figure 5K).**

**Average VAMP content in VGLUT2+ synapses on TA Fast Blue motoneurons** Kruskal-Wallis test (non-parametric):

VAMP1: **p = 0.0034**; K-S Statistic = 11.35

VAMP2: **p = 0.0194**; K-S Statistic = 7.881

Experimental groups:

- Control: non-injected TA,
- TeTx (-): TA injected with TeTx but does not show tetanization,
- TeTx (+): TA injected with TeTx showing positive tetanization.

| Dunn’s multiple comparisons test | mean rank 1 | mean rank 2 | Rank  difference | Mean 1 ± S.D. | Mean 2 ± S.D. | % difference (2-1) | Hedge’s  g | P Value | N1 | N2 | Z |
| --- | --- | --- | --- | --- | --- | --- | --- | --- | --- | --- | --- |
| **VAMP1** |  |  |  |  |  |  |  |  |  |  |  |
| Control vs TeTx (-) | 26.87 | 36.12 | -9.252 | 1,241.0 ± 697.9 | 1,676.0 ± 529.6 | 35.05% | 0.6823 | 0.1781 | 26 | 17 | 1.886 |
| Control vs TeTx (+) | 26.87 | 15.68 | 11.18 | 1,241.0 ± 697.9 | 690.5 ± 598.9 | -44.36% | 0.8203 | 0.1443 | 26 | 11 | 1.976 |
| TeTx (-) vs TeTx (+) | 36.12 | 15.68 | 20.44 | 1,676.0 ± 529.6 | 690.5 ± 598.9 | **-58.80%** | **1.7684** | **0.0024**** | 17 | 11 | 3.357 |
| **VAMP2** |  |  |  |  |  |  |  |  |  |  |  |
| Control vs TeTx (-) | 23.95 | 31.67 | -7.714 | 955.8 ± 538.4 | 1,252.0 ±508.5 | 30.99% | 0.5628 | 0.3093 | 21 | 15 | 1.630 |
| Control vs TeTx (+) | 23.95 | 16.50 | 7.452 | 955.8 ± 538.4 | 684.8 ± 326.5 | -28.35% | 0.5715 | 0.4239 | 21 | 12 | 1.471 |
| TeTx (-) vs TeTx (+) | 31.67 | 16.50 | 15.17 | 1,252.0 ±508.5 | 684.8 ± 326.5 | **-45.22%** | **1.2954** | **0.0155*** | 15 | 12 | 2.797 |

**Supplementary Table 8 (for Figure 6G).**

**% of VGAT+ synapses on TA Fast Blue motoneurons expressing VAMP1 and/or VAMP2**

Kruskal-Wallis test (non-parametric):

VAMP1: **p <0.0001**; K-S Statistic = 34.97

VAMP2: **p <0.0001** ; K-S Statistic = 19.85

Experimental groups:

- Control: non-injected TA,
- TeTx (-): TA injected with TeTx but does not show tetanization,
- TeTx (+): TA injected with TeTx showing positive tetanization.

| Dunn’s multiple comparisons test | mean rank 1 | mean rank 2 | Rank  difference | Mean 1 ± S.D. | Mean 2 ± S.D. | % difference (2-1) | Hedge’s  g | P Value | N1 | N2 | Z |
| --- | --- | --- | --- | --- | --- | --- | --- | --- | --- | --- | --- |
| **VAMP1** |  |  |  |  |  |  |  |  |  |  |  |
| Control vs TeTx (-) | 23.74 | 24.50 | -0.7609 | 99.52% ± 2.32 | 100% ± 0.00 | 0.48% | 0.2534 | >0.9999 | 23 | 12 | 0.2720 |
| Control vs TeTx (+) | 23.74 | 3.500 | 20.24 | 99.52% ± 2.32 | 66.87% ± 8.50 | **-32.81%** | **7.7464** | **<0.0001****** | 23 | 6 | 5.621 |
| TeTx (-) vs TeTx (+) | 24.50 | 3.500 | 21.00 | 100% ± 0.00 | 66.87% ± 8.50 | **-33.13%** | **6.9723** | **<0.0001****** | 12 | 6 | 5.347 |
| **VAMP2** |  |  |  |  |  |  |  |  |  |  |  |
| Control vs TeTx (-) | 25.34 | 28.32 | -2.981 | 84.10% ± 11.11 | 87.31% ± 10.45 | 3.82% | 0.2959 | >0.9999 | 22 | 14 | 0.6820 |
| Control vs TeTx (+) | 25.34 | 4.500 | 20.84 | 84.10% ± 11.11 | 24.49% ± 10.47 | **-70.88%** | **5.4421** | **0.0002***** | 22 | 8 | 3.949 |
| TeTx (-) vs TeTx (+) | 28.32 | 4.500 | 23.82 | 87.31% ± 10.45 | 24.49% ± 10.47 | **-71.95%** | **6.0075** | **<0.0001****** | 14 | 8 | 4.205 |

**Supplementary Table 9 (for Figure 6H).**

**Average VAMP content in VGAT+ synapses on TA Fast Blue motoneurons**

Kruskal-Wallis test (non-parametric):

VAMP1: **p = 0.0185**; K-S Statistic = 7.985

VAMP2: **p = 0.0010**; K-S Statistic = 13.83

Experimental groups:

- Control: non-injected TA,
- TeTx (-): TA injected with TeTx but does not show tetanization,
- TeTx (+): TA injected with TeTx showing positive tetanization.

| Dunn’s multiple comparisons test | mean rank 1 | mean rank 2 | Rank  difference | Mean 1 ± S.D. | Mean 2 ± S.D. | % difference (2-1) | Hedge’s  g | P Value | N1 | N2 | Z |
| --- | --- | --- | --- | --- | --- | --- | --- | --- | --- | --- | --- |
| **VAMP1** |  |  |  |  |  |  |  |  |  |  |  |
| Control vs TeTx (-) | 22.65 | 24.17 | -1.514 | 1,266.0 ± 552.8 | 1,328.2 ± 749.2 | 4.90% | 0.0992 | >0.9999 | 23 | 12 | 0.3550 |
| Control vs TeTx (+) | 22.65 | 8.333 | 14.32 | 1,266.0 ± 552.8 | 517.3  ± 289.6 | **-59.14%** | **1.4557** | **0.0274*** | 23 | 6 | 2.607 |
| TeTx (-) vs TeTx (+) | 24.17 | 8.333 | 15.83 | 1,328.2 ± 749.2 | 517.3  ± 289.6 | **-61.05%** | **1.2629** | **0.0246*** | 12 | 6 | 2.643 |
| **VAMP2** |  |  |  |  |  |  |  |  |  |  |  |
| Control vs TeTx (-) | 26.27 | 25.29 | 0.9870 | 1,094.0 ± 563.6 | 1,070.0 ±621.9 | -2.19% | 0.0443 | >0.9999 | 22 | 14 | 0.2248 |
| Control vs TeTx (+) | 26.27 | 7.250 | 19.02 | 1,094.0 ± 563.6 | 356.3 ± 72.06 | **-67.43%** | **1.5073** | **0.0010**** | 22 | 8 | 3.587 |
| TeTx (-) vs TeTx (+) | 25.29 | 7.250 | 18.04 | 1,070.0 ±621.9 | 356.3 ± 72.06 | **-66.70%** | **1.4183** | **0.0046**** | 14 | 8 | 3.168 |

**Supplementary Table 10**

**Data structure of synaptic samples for the analysis of VAMP1 and VAMP2 content.**

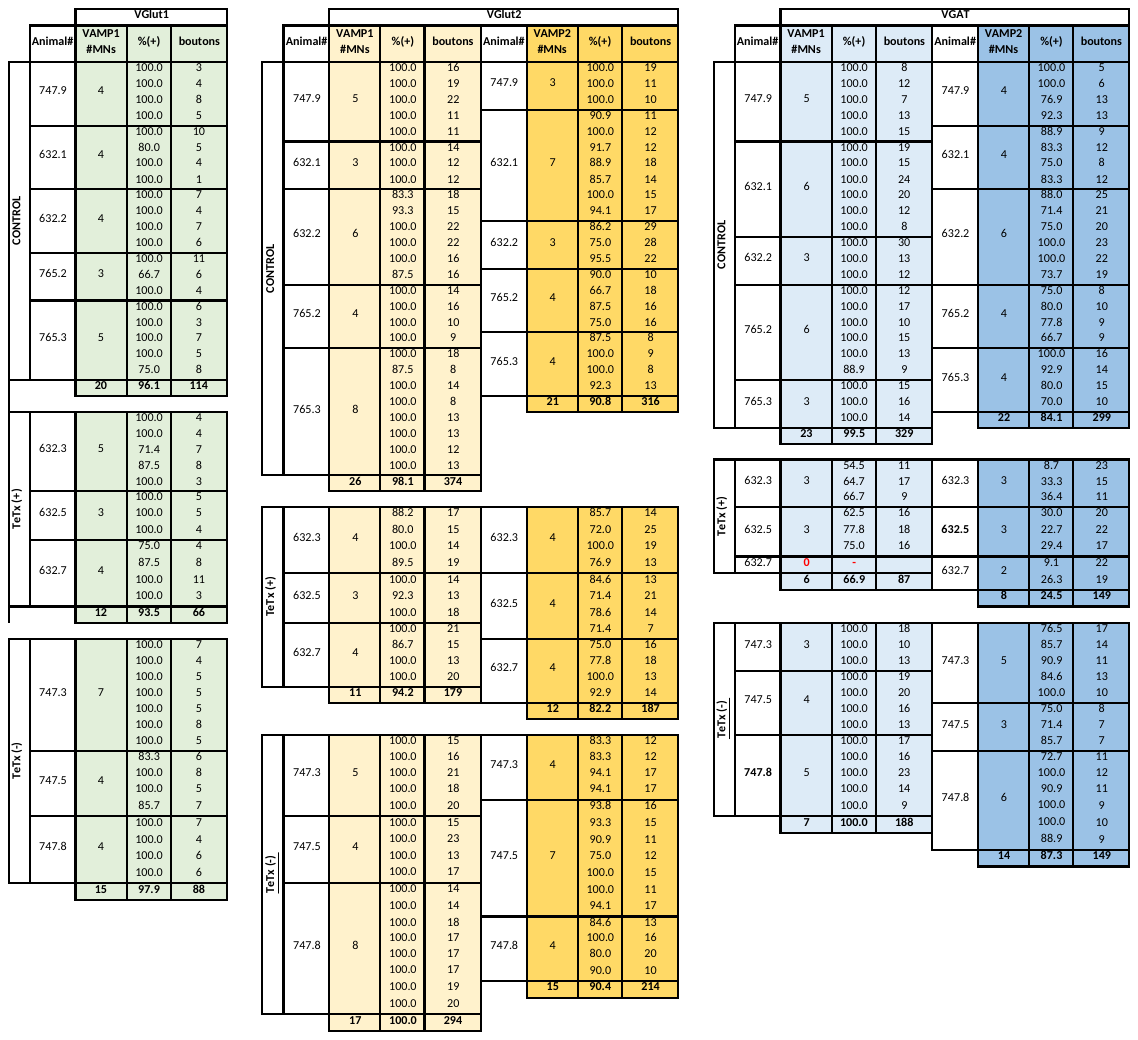
**Supplementary Figure 1**
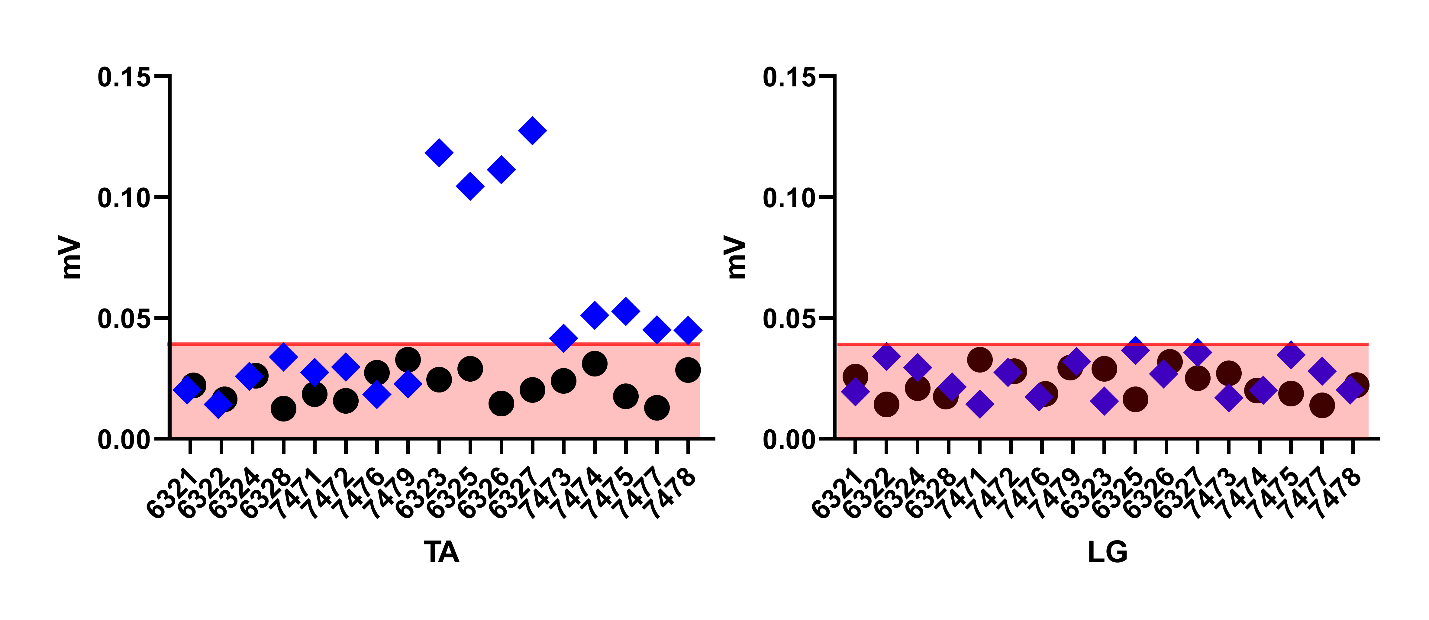

**EMG responses to pressure probe testing in the injected TA and ipsilateral LG muscle** (see text for methods). Outlined circles are data from animals used for analyses of VAMP content. Black circles indicate control side in which the TA was injected with saline (vehicle). Blue rhomboids indicate the side injected with TeTx. Gray band indicates the area containing EMG values within the average +2 SD in TA or LG in control. TA shows positive tetanization when the average EMG response to the pressure probe is above this threshold. Tetanization affected only the injected TA and not the ipsilateral LG. The x-axis represents the animal number (as in Supplementary Table 10). Each animal was tested in the TA and LG of the injected (rhomboid) and non-injected legs (black circles). An additional two animals with no TeTX injections were added to increase motoneurons sampling number and these were not tested with the pressure probe.
